# Single-Molecule Imaging Reveals Transcription-Driven Supercoiling in Unconstrained DNA

**DOI:** 10.1101/2025.11.08.687369

**Authors:** Lili Yang, Yanran Wang, Wei Lyu, Robert G. Egbert, Enoch Yeung

## Abstract

*In vitro* studies of supercoiling dynamics have relied on externally applied force to twist constrained DNA. It is thus unknown whether transcription alone can generate supercoiling in topologically unconstrained DNA, and whether RNAP complexes could act as the topological barriers required to confine this stress. Here, using single-molecule imaging of 20k base pair long double strand DNA, we reveal that RNAP dynamically generates and confines supercoiling in unconstrained DNA. We demonstrate that multiple transcription events create transient topological domains, which allow plectonemes to stabilize between them; in contrast, a single transcription event does not lead to plectoneme formation. Furthermore, we show that this transcription-induced supercoiling is modulated by topoisomerase activity. Crucially, by observing that RNAP itself provides sufficient topological barriers, we establish that transcription-induced supercoiling is an inherently localized phenomenon. These findings redefine the physical basis of transcription-coupled supercoiling, providing a new mechanistic framework for modeling gene regulation based on local topology, with broader implications for genome editing and genome evolution.

## Main

Plectonemes, also known as supercoiled structures of DNA, form in response to torsional stress during various cellular processes^1^. Genomic DNA is frequently subjected to such torsional stress during essential events like transcription^2–4^, replication^5^, recombination^6,7^, and DNA repair^8^. In its topologically relaxed state, DNA adopts a right-handed double-helical conformation with approximately10.5 base pairs per turn^9^. However, when proteins like RNA polymerase (RNAP) translocate along the DNA with minimal rotation, they introduce torsional stress^3,4,10^. This alters DNA twist, resulting in either over-wound or under-wound domains, both of which deviate from the relaxed helical pitch^2^. In topologically constrained systems, such as circular DNA or chromatin, this strain cannot dissipate freely and is instead converted into DNA writhe, forming compact supercoiled structures^4,11–13^. The twin-supercoiled-domain model was formulated to provide a theoretical understanding of this supercoiling process during transcription^2^. It describes the formation of positive supercoils ahead of the RNAP transcription bubble and negative supercoils behind it.

The accumulation of torsional stress and the formation of DNA supercoiling can influence transcription either directly by impeding RNAP progression^3^, or indirectly by altering the landscape of DNA-protein interactions, such as those between the promoter sequence and RNAP itself. To prevent transcriptional stalling and ensure gene expression proceeds normally, cells rely on the timely resolution of this stress either through rotational relaxation of DNA breakage^14,15^ or the activity of topoisomerases^16–19^. Topoisomerases are a family of proteins that relieve supercoiling by transiently cleaving and resealing DNA strands. In *Escherichia coli (E. coli)*, for example, DNA gyrase^20,21^ and topoisomerase I (TOPO-I)^22^ act as a counterbalancing pair to manage positive and negative supercoiling, respectively.

Interestingly, the orientation and transcriptional strength of one gene can influence the expression of neighboring genes^23–25^, suggesting that transcription-generated supercoiling may play a role in modulating local transcriptional activities. To manage this torsional stress, cells have evolved mechanisms to confine supercoiling within defined topological domains. In eukaryotes, DNA supercoiling is largely constrained within the chromatin domains due to nucleosomal organization^5,26,27^. Nucleosomes not only help package DNA but also act as barriers to polymerases and other enzymes, effectively compartmentalizing torsional stress^5,27^. Similarly, in bacteria, abundant nucleoid-associated proteins—such as HU, H-NS, and FIS—aid in DNA compaction and in restraining torsional stress^4,12,28^.

However, the mechanisms underlying these dynamics—particularly whether and how transcription-induced supercoiling by RNAP propagates—remain poorly understood. Previous studies, such as those by Dekker and colleagues, have provided key insights by showing that chemically introduced supercoils become stabilized upon RNAP binding^29^. However, this leaves critical open questions regarding supercoiling that is intrinsically generated by the transcription process itself. These include: While DNA-binding proteins are known to confine supercoiling, do adjacent RNAP complexes play a comparable role in managing torsional stress? In particular, does transcription induce measurable supercoiling in DNA molecules that are unconstrained? To address these questions directly, we visualized the temporal dynamics of transcription-induced supercoiling at the single-molecule level using an array of horizontally stretched unconstrained DNA molecules.

Magnetic tweezers are particularly well-suited for single-molecule studies due to their simplicity, robustness, and versatility^30–36^. This technique enables precise control and measurement of the mechanical properties of DNA, as well as real-time observation of DNA–protein interactions^8,10^ and protein folding^37^. The construction of most magnetic tweezers position a magnetic field source above a flow cell, where individual DNA molecules are tethered to the surface of the coverslip that forms the base of the flow cell^38^.

This configuration allows for the application of torsional and stretching forces to single DNA molecules, while measuring end-to-end distance as an indirect readout of plectoneme size. However, conventional vertical magnetic tweezers cannot directly visualize plectonemes along the DNA molecule in a spatially resolved manner. Although recent modifications have enabled the horizontal visualization of plectoneme structures^31,39^, several challenges persist. These include limited DNA length, difficulties in mimicking physiological transcription conditions, and complex apparatus and sample preparation.

### Direct visualization using horizontal magnetic tweezers reveals transcription-induced plectoneme formation in single DNA molecules

To overcome limitations in studying DNA supercoiling and directly visualize plectoneme formation dynamics along single DNA molecules, we developed a customized horizontal magnetic tweezers apparatus (Fig. 1a). This setup allows Cyanine5 (Cy5)-labeled DNA molecules to be stretched horizontally using either paramagnetic beads or a constant flow through the microfluidic channel of µ-Slide^®^ (Fig. S1). DNA conformational dynamics were visualized using inverted fluorescence microscopy. For each experiment, 20 kb length DNA molecules were PCR-amplified from our in-house plasmid pYW1 using primers modified with either biotin or digoxigenin (DIG). This generated linear DNA molecules with a DIG-modified 5′ end on one strand and a biotin-modified 5′ end on the complementary strand. The DIG-labeled end was anchored to the surface of a µ-Slide^®^ using anti-DIG antibodies, while the biotin-labeled end was attached to a streptavidin-coated paramagnetic bead. The resulting construct was a topologically unconstrained double-stranded DNA molecule, singly tethered not only to the surface but also to the bead.

**Figure 1.**
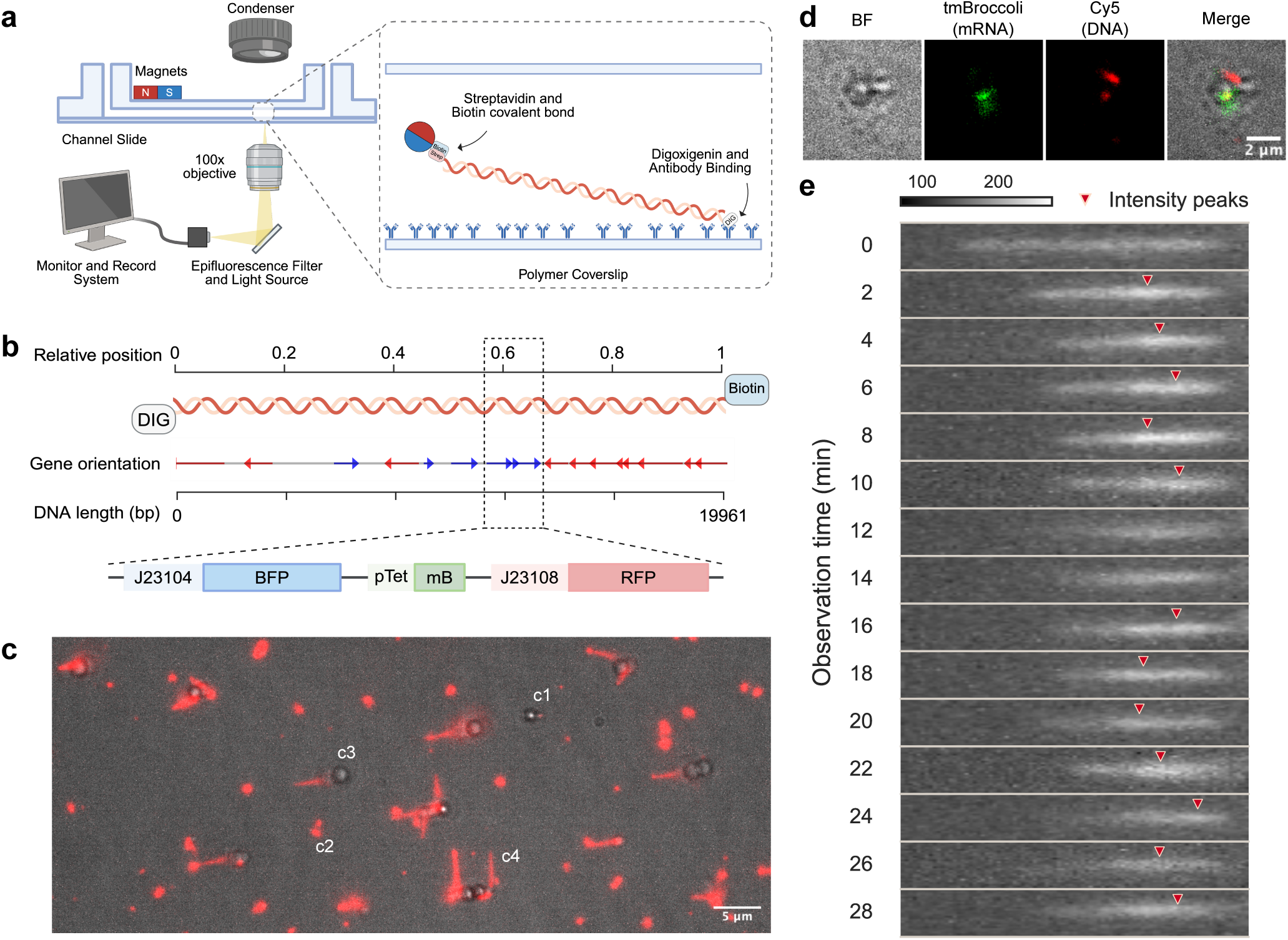
Real-time tracking of transcription-induced supercoiling in unconstrained DNA. **(a)** Schematic of the experimental setup. A 20 kb DNA molecule is asymmetrically labeled with digoxigenin (DIG) at the 5′ end of one strand and biotin at the 5′ end of the complementary strand. The DIG-labeled end is tethered to the surface of a µ-Slide that coated with anti-DIG, while the biotin-labeled end is attached to a streptavidin-coated paramagnetic bead. The DNA molecule is stretched horizontally by a stack of magnets positioned atop and alongside of the flow channel. DNA conformations are monitored in real time using an inverted fluorescence microscope. **(b)** Schematic of the 20 kb DNA construct, mapped to a normalized [0, 1] scale for visualization. The sequence is derived from the *E. coli* genome and includes three synthetic promoters (all in the same orientation) driving fluorescent reporters near the 0.6 position. mB: tmBroccoli RNA aptamer. **(c)** Representative composite image of a tethered DNA molecule under a magnetic field, showing bright-field (gray) and Cy5-labeled DNA (red). The observation was performed at 37°C in transcription buffer. Scale bar: 5 µm. **(d)** Visualization of DNA and synthesized tmBroccoli mRNA in transcription buffer supplemented with RNAP and DFHBI fluorophore, after a 45-minute incubation at 37°C. Images from left to right: bright-field, tmBroccoli fluorescence (mRNA), Cy5 fluorescence (DNA), and a merged composite. Scale bar: 2 µm. **(e)** Real-time analysis of DNA conformational changes in the presence of transcription buffer and RNAP at 37°C, presented as a time-series image marked with fluorescence intensity peaks. The illustration figures (a and b) were created in BioRender (https://BioRender.com).

The linear DNA consists mainly of *E. coli* genomic DNA and a 1.9 kb synthetic insert (sequence available in Supplementary Material and Method). The insert features a tetracycline-responsive promoter (pTet) regulated tmBroccoli^40^ RNA aptamer gene and two constitutively expressed fluorescent protein genes, situated approximately 8 kb from the biotin-labeled end (Fig. 1b). When imaged, most beads were properly tethered to DNA molecules. Nevertheless, some beads adhered directly to the µ-Slide surface (Fig. 1c1), while some DNA molecules lacked bead attachment at the biotinylated end (Fig. 1c2). Under the magnetic field, most DNA molecules with bead attached were stretched in line with the field direction (Fig. 1c3). However, when multiple DNA-bead complexes were in close proximity, the beads attract one another, aligning into linear aggregates along the magnetic field. This caused some of the stretched DNA molecules to adopt non-uniform orientations (Fig. 1c4). The average length of stretched DNA under magnetic force was approximately 3.47 ± 0.79 µm (Fig. S2).

Before investigating transcription-induced supercoiling, we confirmed that Cy5 labeling did not interfere with transcriptional activity. We verified this using two complementary approaches: single-molecule microscopy and *In vitro* plate reader assays. At the single-molecule level, transcription proceeded normally, as measured by tmBroccoli synthesis after a 45-minute incubation under the microscope (Fig. 1d). In plate reader assay, the transcription capability was further confirmed by collecting time series data of tmBroccoli expression up to 12 hours (Fig. S3). To visualize transcription-induced conformational changes of DNA, RNAP was added to the µ-Slide^®^ channel to initiate transcription on tethered DNA molecules. To capture the resulting dynamics, we performed time-lapse imaging at 2-minute intervals with a 200-millisecond exposure per frame. We observed that plectonemes appeared as Cy5 signal peaks on the DNA kymograph after just 2 minutes of incubation with RNAP at 37 °C (Fig. 1e).

Our customized horizontal magnetic tweezers apparatus thus provides a powerful tool for dissecting the real-time conformational dynamics of DNA under transcription-induced torsional stress (a schematic overview of the experimental and analytical workflow shown in Fig. S4). This system permits detailed *in vitro* investigations into the kinetics of plectoneme formation, their stability, and their interplay with topoisomerases during active transcription.

### RNAP activity-dependent transcription drives DNA conformation dynamics

To determine whether DNA conformational change observed is caused by RNAP activity-dependent transcription process, DNA conformational dynamics were analyzed over time at different RNAP activity levels. RNAP activity is governed by multiple factors throughout a series of well-defined stages^41^. First, RNAP binds to promoter region^42^ and form an open complex, allowing local DNA unwinding^43,44^. Formation of the open complex is more thermodynamically favorable at 37 °C than at lower temperatures^45,46^. Transcription then begins with abortive initiation, a process catalyzed by the β subunit of RNAP; the short RNA transcripts produced are also released through a channel in the β subunit. When the β subunit is blocked by an inhibitor such as rifampicin, RNAP becomes trapped in the abortive initiation cycle and cannot transition to the elongation phase, although its binding to DNA is not impaired^47,48^. Finally, RNAP synthesizes the mRNA by incorporating nucleotides during elongation, a process that follows the Q₁₀ rule, whereby catalytic activity approximately doubles with a 10 °C increase in temperature^49,50^. Therefore, we employed a temperature series (29°C, 33°C and 37°C) and rifampicin inhibition to establish a range of RNAP activity levels. Consequently, transcriptional activity of tmBroccoli showed a significant reduction at 29 °C and 33 °C than at 37 °C (Fig. S5). Moreover, rifampicin treatment effectively prevented tmBroccoli synthesis as expected (Fig. S6).

In our analysis of DNA conformation, we found that the DNA molecules underwent a phase transition from linear to condensed, and finally to detached state (Fig. S7). We defined condensed DNA as a compact structure with a diameter less than 1.2 µm, a limit set by microscopic spatial resolution. Time-lapse image analysis revealed that under optimal RNAP activity at 37 °C, linear DNA molecules progressed through condensation to detachment at a significantly accelerated rate compared to conditions with impaired RNAP activity—for example, at lower temperatures or with rifampicin inhibition (Fig. S7 and S8). These results suggest that the phase transition from condensation to detachment was attenuated at lower RNAP activities. Specifically, the overall DNA shrinkage observed throughout the experiment was most pronounced in the active RNAP (37°C) group. This difference was statistically significant (p < 0.001) when compared to all other conditions, including the no-RNAP control and groups with RNAP at 29°C, 33°C, or with rifampicin (Fig. 2a). This result directly links the magnitude of DNA shrinkage to the level of transcriptional activity.

**Figure 2.**
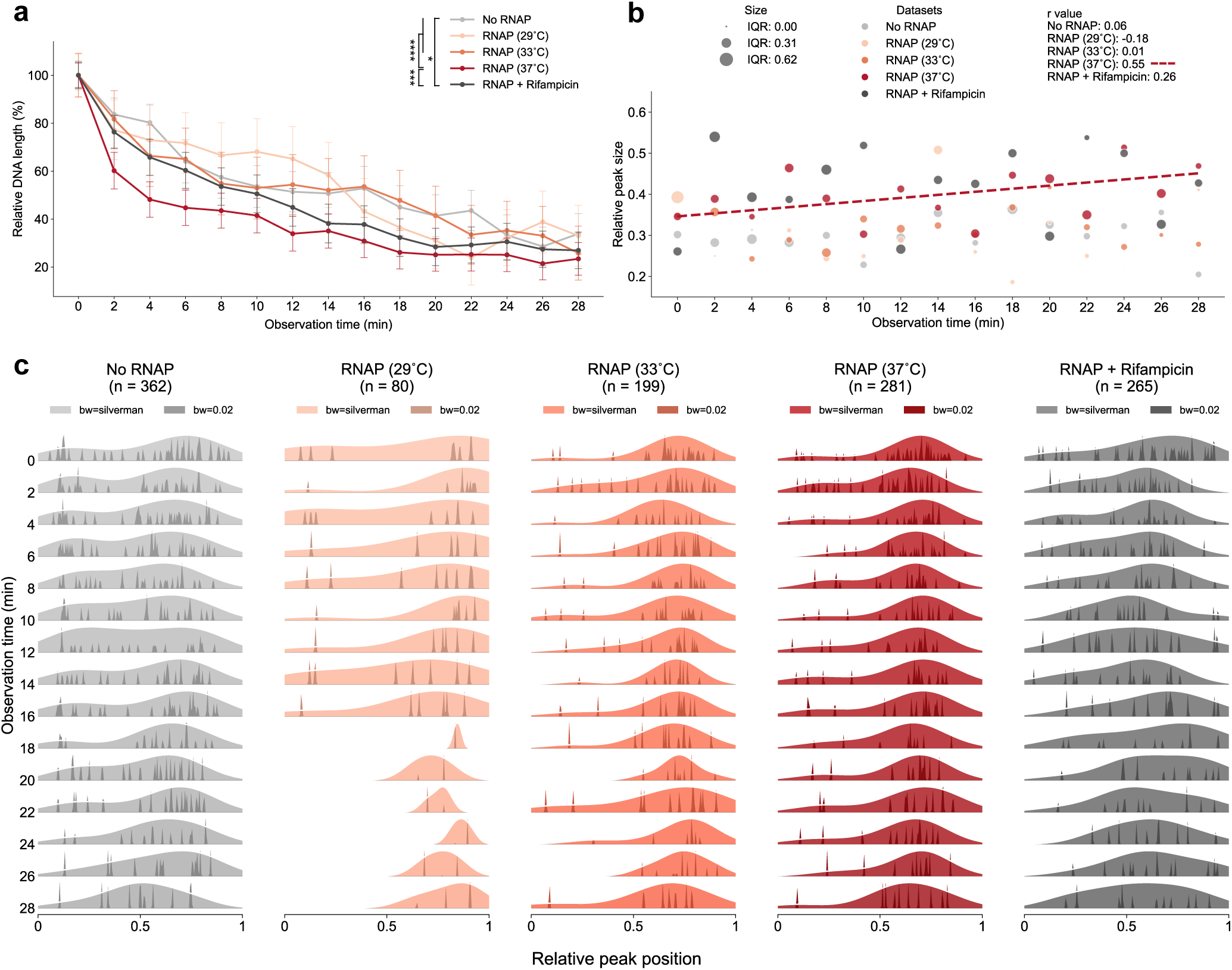
Time-resolved DNA dynamics reveal RNAP activity-dependent supercoiling. **(a)** DNA length changes over time, normalized to the initial length. Data are presented as mean ± standard error of the deviation (sed). Statistical comparisons were performed using a mixed-effects model followed by Tukey’s HSD post-hoc test (*p < 0.05; ***p < 0.001; ****p < 0.0001). The number of DNA molecules analyzed for each experimental group and time point were as follows: No RNAP (N = 48), RNAP at 29°C (N = 17), RNAP at 33°C (N = 26), RNAP at 37°C (N = 60), and RNAP with Rifampicin (N = 47). **(b)** DNA peak sizes, normalized to DNA length, tracked over time. Pearson’s correlation coefficient (r) was calculated for each group. A dashed line indicates the linear regression for groups where |r| > 0.5. The interquartile range (IQR) is shown for each time point. **(c)** DNA peak positions plotted on a normalized [0, 1] scale. For each frame, peak position distributions were estimated using Gaussian kernel density estimation (KDE) with Silverman’s bandwidth (bw) method. Density values were normalized to half of their maximum for visual clarity. Conditions: No RNAP: DNA with transcription buffers only. 29°C, 33°C, and 37°C: DNA with transcription buffer and RNAP at the indicated temperatures. RNAP + Rifampicin: the *E. coli* RNAP inhibitor was supplemented in transcription buffer at a final concentration of 1 µg/mL. The value of *n* represents the total number of intensity peaks analyzed across all time points for each experimental group.

### RNAP activity governs the spatial distribution of transcription-induced supercoils

Plectonemes appear as distinct region due to their compacted DNA structure, which exhibit higher fluorescence intensity than relaxed segments (Fig. S9). Despite its relevance, the behavior of plectonemes formed during transcription is still not well characterized. We analyzed the spatial distribution of plectonemes through time by identifying localized intensity peaks along all DNA molecules in each group. In all the groups, peak numbers that identified ranges from 80 to 362 peaks. The identified peaks were plotted on a relative scale of DNA molecule ranges from 0 to 1. Meanwhile, the numbers of peaks at each relative position were plot as the high on y-axis. Finally, the smoothed curve of plectoneme positions were presented as probability density (Fig. 2b).

Without RNAP presence, the probability density peaks showed temporal instability, frequently reverting to a more uniform distribution (indicating no persistent plectonemes). This contrasts sharply with RNAP-treated DNA molecules at 37°C, which maintained stable, defined peaks throughout observations. Under active transcription, the probability density of plectoneme positions formed a left-skewed distribution peaking at a relative position of 0.7, reflecting consistent positioning preferences across DNA molecules. This emergent pattern shows clear transcription-dependent stabilization of plectonemes at specific loci. Rifampicin treatment disrupted this pattern, eliminating the stable positioning and further confirming its dependence on transcriptional activity.

The effect of temperature was primarily on the rate of plectoneme formation, rather than on the spatial distribution of the plectonemes themselves. At 29°C, the appearance of stable plectoneme peaks was significantly delayed, first emerging around the 20-minute mark. At 33°C, the delay was less pronounced, with peaks first appearing around 8 minutes. However, once established at both temperatures, the peaks stabilized temporally, and their positions were identical to those observed at 37°C. This indicates that while temperature influences the rate of supercoiling emergence, it does not alter the spatial distribution of the stable plectonemes. The predominant peak position near 0.7 observed in the RNAP-treated group corresponds to the region of the inserted genes, which are driven by stronger promoters than those of the surrounding genes. This suggests that plectoneme formation are strongly influenced by transcriptional activity and promoter strength.

When restricting the analysis to intensity-profile peaks (plectonemes) occurring between relative positions 0.1 and 0.9 to exclude boundary noise, we observed a linear increase in plectoneme size over time with RNAP presence at 37 °C, ranging from ∼6 kb to ∼9 kb. In contrast, the other four groups, which either lacked RNAP or had attenuated RNAP activity, showed minimal temporal correlation in plectoneme size (Fig. 2c).

Supported by transcriptional activity data (Fig. S5), these results collectively demonstrate that the observed peaks represent DNA supercoiling directly induced by RNAP transcription. Furthermore, the formation of these supercoils exhibited distinct temporal dynamics for each group. Both the rate of plectoneme emergence and plectoneme size showed a positive correlation with RNAP activity. In contrast, the spatial distribution of plectoneme along the DNA molecule remained largely conserved across different levels of RNAP activity.

### DNA gyrase and TOPO I introduce oscillatory dynamics in DNA supercoiling

Next, we investigated how DNA gyrase and TOPO I influence DNA supercoiling. To establish an optimal condition for single-molecule analysis, we first needed to determine effective enzyme concentrations. We used a plate reader to monitor tmBroccoli expression over 12 hours in transcription buffer, reasoning that the combination causing the largest deviation in transcription from the RNAP-only control would indicate the strongest perturbation of supercoiling dynamics. All tested gyrase:TOPO I ratios significantly altered the transcriptional outcome (Fig. S10). The 0.2:1 gyrase:TOPO I ratio produced the most significant deviation, with a final signal 65% greater than the RNAP-only control— substantially exceeding the 39% and 53% differences of the 0.1:1 and 1:1 ratios, respectively. We therefore selected the 0.2:1 ratio for subsequent single-molecule experiments to maximize the observable effect.

In the presence of gyrase and TOPO I, single-molecule imaging revealed an oscillating pattern in DNA length over time, contrasting with the linear decrease observed in the RNAP-only group (Fig. 3a). Notably, statistical DNA length analysis also demonstrated oscillatory patterns (Fig. 3b), confirming that the phenomenon is statistically consistent across molecules. These dynamics reflect competing mechanical influences on DNA length (Fig. 3c): (1) magnetic field stretching forces that extend the DNA, (2) transcription-induced compaction that opposes extension, and (3) topoisomerase-mediated length modulation that regulates the two forces above. We further calculated and fitted the applied force on DNA molecules using the Marko–Siggia worm-like chain (WLC) approximation^51^ (Fig. S11). While these “forces” represent theoretical constructs derived from length dynamics rather than direct measurements, their oscillatory behavior demonstrates coordinated topological regulation. Specifically, time-resolved comparisons between RNAP-only and gyrase-TOPO I groups show that topoisomerases periodically relieve torsional stress (Fig. S12), while the persistent negative correlation between transcription and magnetic forces confirms their opposing mechanical roles in shaping DNA length (Fig. 1e).

**Figure 3.**
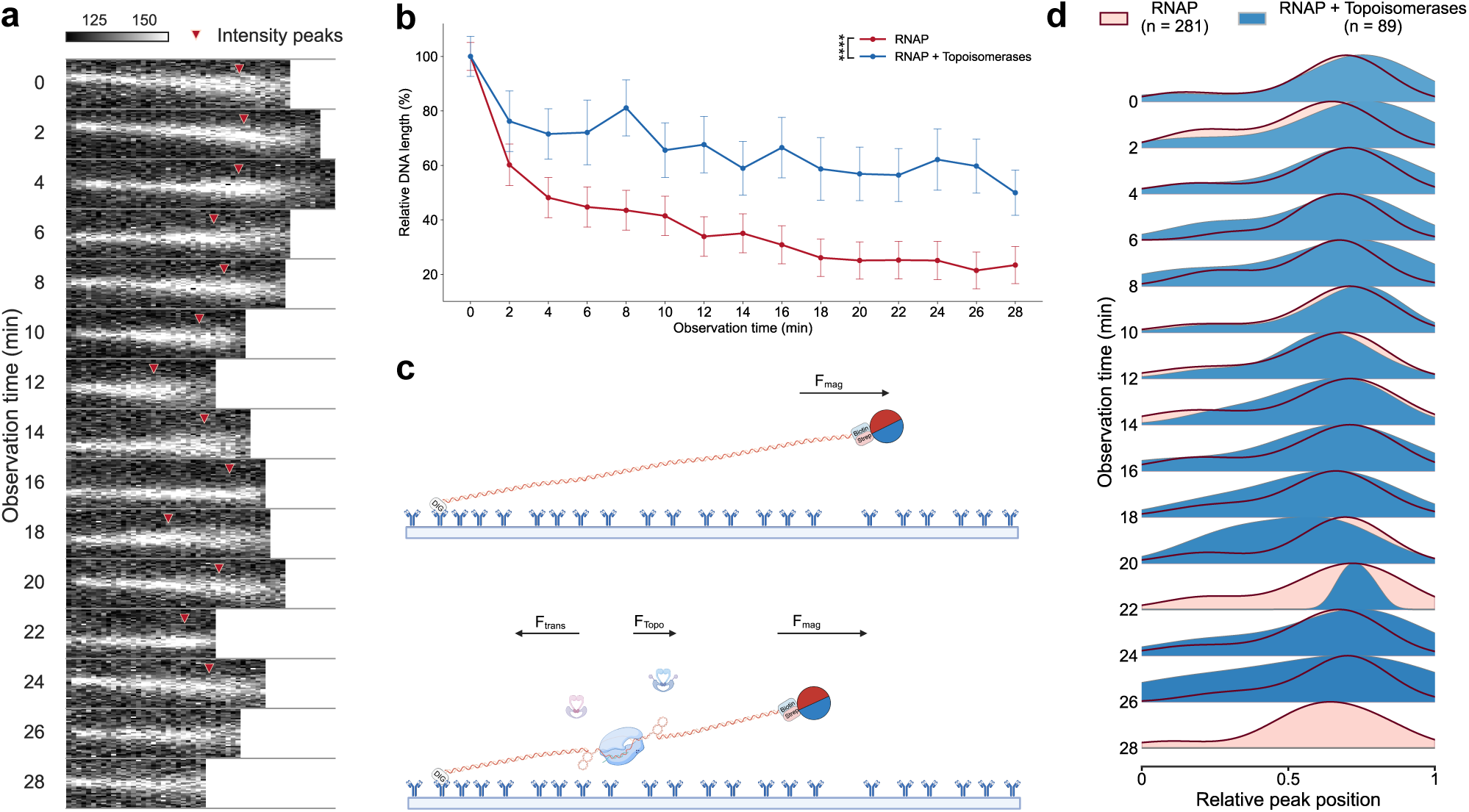
Gyrase and Topo I relieve transcription-induced torsional stress and induce oscillatory DNA conformation dynamics. **(a)** Real-time analysis of DNA conformational changes in transcription buffer supplemented with RNAP, and a combination of gyrase and topoisomerase I at 0.2:1 ratio, presented as a time series image marked with fluorescence intensity peaks. **(b)** Schematic illustrating the forces acting on the DNA: the magnetic stretching force (F_mag_) that extends the DNA, the transcription-induced compaction force (F_trans_) that opposes extension, and the topoisomerase-mediated force that relieves torsional stress (F_topo_). This illustration figure was created in BioRender. **(c)** DNA length changes over time, normalized to the initial length. Data are presented as mean ± standard error of the deviation (sed). Statistical comparisons were performed using a mixed-effects model followed by Tukey’s HSD post-hoc test (****p < 0.0001). The number of DNA molecules analyzed for each experimental group and time point were as follows: RNAP (N = 60), and RNAP with Topoisomerases (N = 8). **(d)** Distribution of DNA intensity peak positions plotted on a normalized [0,1] scale. For each frame, peak position distributions from the two experimental groups are overlaid. Peak position distributions were estimated using Gaussian kernel density estimation (KDE) with Silverman’s bandwidth (bw) method. Density values were normalized to half of their maximum for visual clarity. The absence of a blue ridgeline at the 28-minute timepoint in the RNAP + Topoisomerases group indicates that fewer than two peaks were identified. The value of *n* represents the total number of intensity peaks analyzed across all time points for each experimental group.

This periodic release of torsional stress distinctly affected plectoneme dynamics. The spatial positions of plectonemes remained relatively stable over time and were nearly identical to those in the RNAP-only group, indicating that topoisomerase activity (specifically of gyrase and TOPO I) does not alter plectoneme locations (Fig. 3d). In contrast, plectoneme size exhibited oscillatory dynamics, demonstrating that topoisomerases periodically release torsional stress to modulate plectoneme magnitude (Fig. S13).

These findings demonstrate that the combined activity of DNA gyrase and TOPO I modulates DNA length *in vitro* and supercoiling magnitude while maintaining the spatial positioning of supercoils. This regulatory dynamic provides mechanistic support for our conclusion: the observed intensity peaks represent DNA supercoiling, as their behavior is directly governed by topoisomerases.

### Multiple RNAP binding events may contribute to supercoiling on unconstrained DNA

In a manually twisted magnetic tweezer system, supercoils cannot form on a singly tethered DNA fragment because the molecule is topologically unconstrained (Supplementary movie 1 and 2). This finding prompts a key question: why did we observe transcription-induced supercoiling in our system, which also used topologically unconstrained DNA? Could it be that neighboring RNAP complexes play a role in managing DNA torsional stress? We therefore hypothesize that RNAP binding sites and/or transcription bubbles act as transient physical barriers, effectively segmenting the DNA molecule and enabling the accumulation of local supercoiling during transcription. Having already observed this local supercoiling, we next sought direct evidence for the multiple RNAP binding events that underlie this segmentation framework.

To resolve this, we imaged the binding positions of RNAP along individual DNA molecules using RNAP labeled with Alexa 594-conjugated antibodies. A representative image after a 10-minute incubation shows multiple RNAP binding events on a single 20 kb DNA molecule (Fig. 4a). As this pattern was representative of a broader trend, we proceeded to quantify RNAP binding across multiple molecules. The resulting profiles, generated by plotting Alexa 594 fluorescence intensity peak against relative DNA position, confirmed the simultaneous binding of multiple RNAPs was a common occurrence (Fig. 4b). By aligning the observed RNAP binding positions with the known coding sequence of our DNA construct (Fig. 4c), we found that RNAP binding sites preferentially match to gene coding regions. Furthermore, single-molecule time series analysis of RNAP and DNA co-localization revealed that DNA supercoiling tends to accumulate in regions flanked by two RNAP binding regions (Fig. 4d). On the contrary, the Alexa 594 fluorophore showed no binding or accumulation effects on DNA molecules, suggesting those co-localization only happened in RNAP presence (Fig. S14).

**Figure 4.**
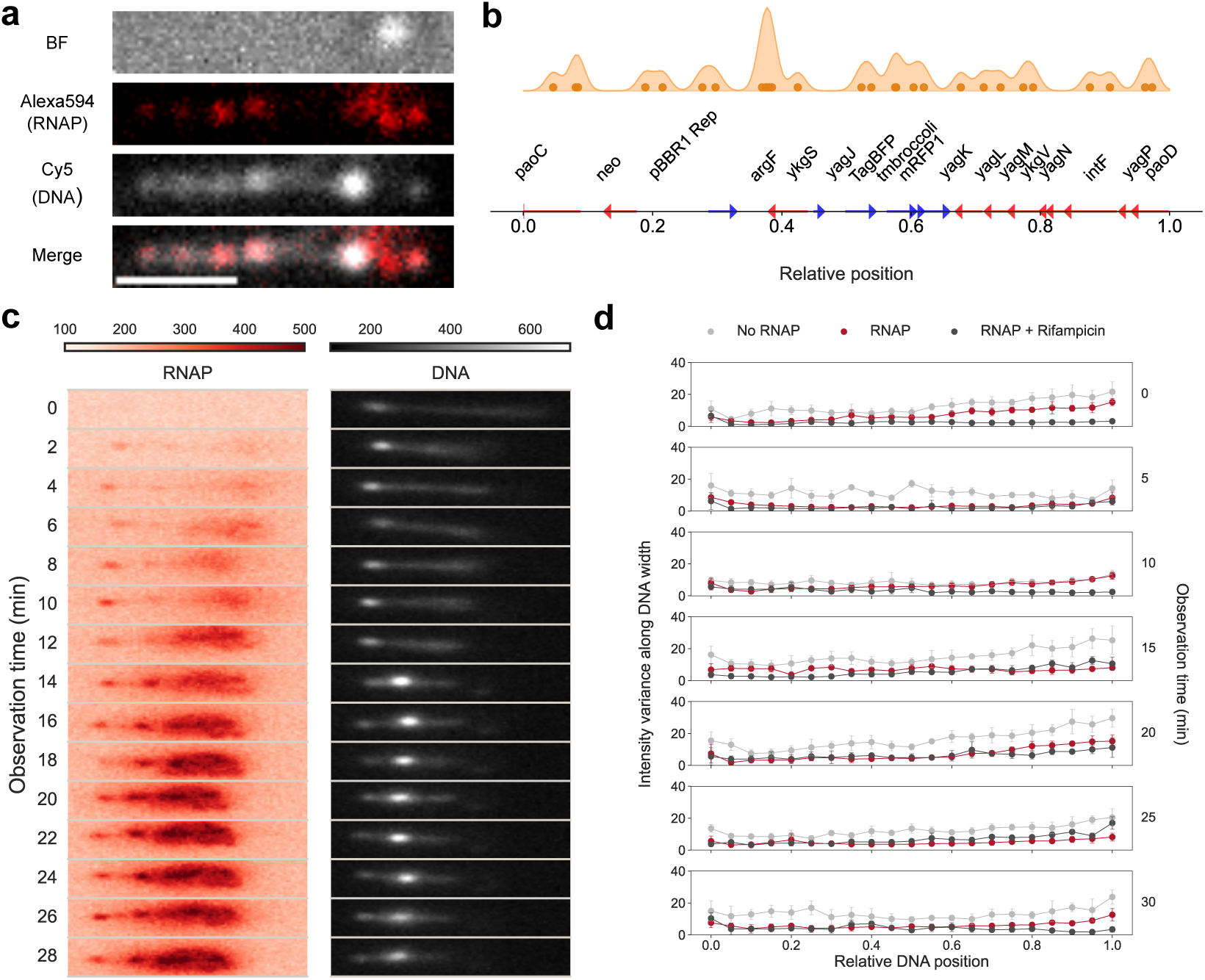
Multiple RNAP binding events mediate transcription-induced torsional stress and enable supercoiling accumulation. **(a)** Representative images showing the co-localization of DNA and RNAP after a 10-minute incubation with transcription buffer supplemented with RNAP, an anti-RNAP primary antibody, and an Alexa Fluor 594-conjugated secondary antibody. Panels from top to bottom: bright-field (gray), Alexa Fluor 594 signal (RNAP, red), Cy5 signal (DNA, black and white), and a merged composite of RNAP and DNA signals. Scale bar: 2 µm. **(b)** Analysis of RNAP binding sites. (Top) Intensity peaks of Alexa Fluor 594-labeled RNAP (n = 8 molecules) plotted against the relative DNA length. (Middle) Gene names and their relative positions. (Bottom) Gene schematic showing orientation and size along the relative DNA scale. **(c)** Real-time analysis of RNAP–DNA co-localization dynamics. Time series images of Alexa Fluor 594-labeled RNAP (left) and Cy5-labeled DNA (right). **(d)** Intensity variance along the y-displacement versus x-position (relative DNA length). For each group, 30 consecutive frames were captured at each timepoint with a 200-millisecond exposure time. For each experimental group, six of DNA molecules were analyzed.

In addition, we recorded DNA molecule vibrations at seven timepoints at 5-minute intervals. At each timepoint, DNA was imaged consecutively for 6 seconds. We analyzed the vertical variance of fluorescence intensity along the DNA. For DNA in transcription buffer alone, significant fluctuations were observed at all timepoints, reflecting consistent active thermal motion at 37°C. In contrast, in the presence of RNAP, DNA molecules exhibited reduced fluctuations and greater overall stability over time, suggesting that RNAP binding suppressed DNA vibration (Fig. 4e and supplementary movie 3). A similar pattern was observed when RNAP activity was inhibited by rifampicin, indicating that the RNAP remained bound and stabilized the DNA even after transcriptional inactivation (Fig. S6 and Fig. 4e).

In this section, we obtained the direct evidence we sought for multiple, simultaneous RNAP binding events on individual DNA molecules. This finding confirms the foundational premise of our segmentation framework, providing a mechanistic explanation for how transcription-induced supercoiling can emerge and be confined within a topologically unconstrained DNA system.

### Transcription events on neighboring genes trap DNA supercoiling in between

Our findings provide clear answers to the central questions posed at the outset of this study. We demonstrate that adjacent RNAP complexes do indeed play a comparable role to other DNA-binding proteins in managing torsional stress, not merely as generators but as crucial confiners of supercoiling. Specifically, we have shown that transcription does induce measurable supercoiling in topologically unconstrained DNA, precisely because RNAP binding and transcription bubbles themselves define effective boundaries. This segmentation creates local topological domains where supercoils can accumulate, and we have quantified the distinct spatial and temporal dynamics of these plectonemes, showing their positive correlation with RNAP activity (Fig. 2). This mechanistic explanation naturally leads to a critical prediction: a single transcription event, in the absence of multiple RNAPs to form barriers, would allow transcription-induced torsional stress to dissipate freely along the DNA. Consequently, without this confinement, no stable, observable plectoneme could accumulate.

However, a key limitation of our initial system was that the barrier effect of RNAP was experimentally inseparable from the act of transcription itself. To test our prediction and overcome this limitation, we engineered a new construct using T7 promoter—which is exclusively recognized by T7 RNAP (Fig. S15)—to drive the tmBroccoli gene, while retaining the flanking constitutive *E. coli* promoters. When only T7 RNAP was introduced, no DNA supercoiling was observed (Fig. 5a and S16). However, when both T7 RNAP and *E. coli* RNAP were added, DNA supercoiling reappeared and increased in size temporally (Fig. 5a, 5b, S16). This confirms that concurrent transcription from neighboring genes is essential for the net accumulation of torsional stress.

**Figure 5.**
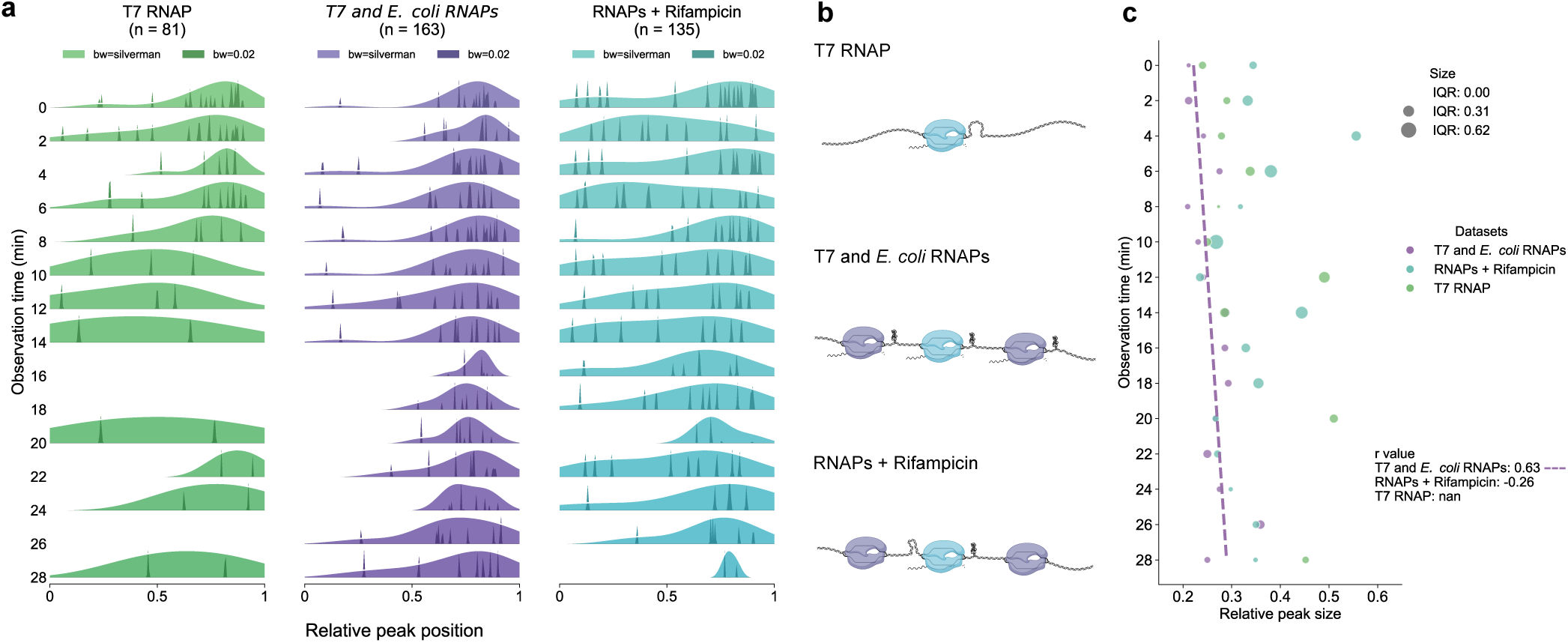
Neighboring transcription events constrain supercoiling in unconstrained DNA. **(a)** DNA signal peak positions plotted on a relative [0, 1] scale. DNA molecules were incubated under three conditions: (1) T7 RNAP only; (2) T7 RNAP with *E. coli* RNAP; and (3) T7 RNAP, *E. coli* RNAP, and the inhibitor rifampicin. The absence of a green ridgeline at the 16 and 26-minute time point in the T7 group indicates that fewer than two peaks were identified. The value of *n* represents the total number of intensity peaks analyzed across all time points for each experimental group. bw: bandwidth. **(b)** Schematic diagram illustrating the proposed model of supercoiling constraint under the three experimental conditions. This illustration figure was created in BioRender. **(c)** DNA peak size normalized to DNA length over time. Pearson’s correlation coefficient (r) was calculated for each group. A dashed line indicates the linear regression for groups where |r| > 0.5. The interquartile range (IQR) is shown for each time point.

To isolate the effect of T7 RNAP, we inactivated *E. coli* RNAP with rifampicin. Under these conditions, T7 RNAP-driven supercoiling exhibited a significant delay of approximately 10 minutes before appearing at relative position near 0.7 (Fig. 5a). This pattern is the inverse of that observed with the *E. coli* promoter-only construct treated with rifampicin (Fig. 2c), where a slight, transient formation of supercoiling was completely disrupted within 10 minutes. The stark contrast confirms that the late appearing plectonemes must result from the gradual accumulation of DNA supercoiling by T7 RNAP initiated transcription.

Fundamentally, the visibility of a plectoneme represents a balance between two opposing dynamics, analogous to a classic rate problem: the production of supercoils by transcription and their dissipation through the surroundings. The delayed appearance of T7-induced supercoiling indicates that the net rate of accumulation is much slower than in a multi-gene system. At this stage, we cannot separate whether this is due to a lower rate of DNA supercoiling generation by a single gene, a faster rate of dissipation due to weaker topological barriers, or a combination of both. Our experiment defines the net effect but cannot yet deconvolve the individual contributions of production and dissipation.

Together, these findings indicate that observable supercoiling in unconstrained DNA is an emergent property of coordinated transcription from multiple genes. To prevent the dissipation of torsional stress, the system requires both the stress generated by transcription itself and the transient physical barriers formed by neighboring transcription complexes.

### Conclusion and discussion

In this work, we showed that transcription generates DNA supercoiling even in topologically unconstrained DNA molecules. Multiple RNAP binding events create localized physical barriers that impede torsional stress dissipation, thereby sustaining supercoiling (Fig 2, Fig 5 and Supplementary movie 3). Intriguingly, topoisomerases (gyrase and TOPO I) resolve this supercoiling in an oscillatory manner, revealing the counteracting dynamics between transcription and topoisomerase activity on DNA supercoiling. These findings not only redefine how transcription regulates DNA topology *in vitro*, but also provide a new framework for theoretical modeling of supercoiling dynamics.

For temporal resolution optimization, we employed a 2-minute imaging interval to minimize photobleaching of Cy5 and Alexa 594 fluorophores. Photobleaching is primarily driven by reactive oxygen species (ROS) generated under high excitation energy, which may cause irreversible damage to fluorophores^52,53^. While oxygen scavenging systems (e.g., oxidase/catalase mixtures) can effectively suppress this effect by reducing ROS production^10,54,55^, their scavenging efficiency is impaired by thiol-containing compounds such as dithiothreitol (DTT)^56,57^. Since DTT is required to maintain RNAP activity, careful optimization of the imaging buffer is essential in balance fluorophore photostability with RNAP functionality. Moreover, the scavenging system^54^ we tested was incompatible with the fluorophore DFHBI used for RNA aptamer labeling, likely due to compound crystallization (data not shown). We are further limited by the effective time window for observation since the majority of detachments occurred within the first 30 minutes (Fig. S7 and S8), which is shorter than that required for tmBroccoli expression quantification (Fig. 1d). As a result of these chemical and temporal constraints, while we were able to visualize supercoiling over time, we could not simultaneously observe real-time RNA synthesis in this study. An improved oxygen scavenging system that enhances photostability while preserving RNAP activity and maintaining RNA labeling compatibility would enable simultaneous visualization of transcription-induced DNA supercoiling and RNA production at sub-minute resolution.

Additionally, due to limited spatial resolution of the microscope and the close spacing of transcribing genes, plectonemes appeared with a size range from 6-9 kb covering 8-10 genes. These appeared as diffraction-limited dots in our observations, which did not allow us to resolve supercoiling at the level of individual genes. According to the theoretical twin-domain model, transcription generates positive supercoiling downstream and negative supercoiling upstream of the transcription bubble^2^. However, the spatial resolution limitations prevent direct visualization of the predicted two different supercoiling domains. To improve spatial resolution, using long DNA templates with single-gene promoters could achieve the isolation of the two supercoiling domains. This approach would enable the precise characterization of transcription-coupled supercoiling dynamics, providing the data necessary to construct a quantitative model from first—the primary focus of our ongoing work.

DNA supercoiling generates certain compaction force in the direction opposite to the applied magnetic field, leading to progressive shortening of DNA molecules over time (Fig. 2c and Fig. 3a). Theoretically, a 20 kb DNA molecule has a relaxed linear length of approximately 6.8 µm^58^ and can compact to 0.14-0.68 µm^33^. In our experiments, the initial lengths of 20 kb DNA molecules ranged from 1.9 to 6.0 µm and consistently shortened to less than 1.2 µm at later timepoints. This observation suggests that RNAP alone is sufficient to drive DNA compaction, potentially forming densely coiled structures or RNAP-supercoil complexes.

Upon relieving supercoiling through topoisomerase treatment (Fig. 3), transcription levels significantly increased (Fig. S10), suggesting that accumulated supercoiling can negatively impact gene expression. DNA gyrase and TOPO I play complementary roles in managing supercoiling stress^16,19^. Gyrase consumes ATP to introduce negative supercoils^20,21,59,60^, whereas TOPO I—frequently associated with RNAP—relaxes them^17,22,61^. Together, they maintain a dynamic balance of supercoiling and relaxation during transcription. Our data support this model, revealing oscillatory patterns in DNA supercoiling and relaxation over time (Fig. 3). Interestingly, variation in the DNA gyrase to TOPO I ratio is likely a key determinant of the oscillations’ amplitude and frequency, as transcription levels were sensitive to changes in this ratio (Fig. S10). Although it is now clear that topoisomerases assist in relieving transcription-induced supercoiling, how their relative concentrations—and specifically their ratio to RNAP—modulate supercoiling dynamics remains an open question. A deeper understanding of how these enzyme ratios influence DNA conformation will be essential for refining predictive models of transcription-coupled supercoiling.

Our findings call for revisions to current models of transcription-coupled supercoiling, particularly in systems involving topologically relaxed or unconstrained DNA. The identification of RNAP as a topological barrier and the oscillatory behavior of topoisomerase activities provide new mechanistic insights into the dynamic regulation of gene expression. This localized interplay between transcription and DNA supercoiling raises broader implications, including concerns about the safety of gene editing. Edited genes may unintentionally alter the transcriptional dynamics of neighboring genes, especially when these genes are not part of the same operon or regulatory cassette—an effect that may be overlooked by current genome recombineering practices. Furthermore, this local physical interplay of supercoiling and transcription suggests that gene transcription strength and genome spatial organization may have co-evolved under the influence of supercoiling dynamics. Thus, transcriptional regulation via DNA topology should be considered a distinct regulatory network—parallel to, but independent of— traditional biochemical signaling pathways. Future research should aim to integrate these physical mechanisms into genome-wide studies and evaluate their relevance in living systems.

## Acknowledgements

We are grateful to Cees Dekker and Terence Strick for insightful discussions. We also acknowledge all members of the Biological Control Lab for their support and for fostering a stimulating research environment. This work was funded in part by an NSF CAREER Award 2240176, the Army Young Investigator Program Award W911NF2010165 and the Institute of Collaborative Biotechnologies/Army Research Office grants W911NF1920026, W911NF19D0001, W911NF22F0005, W911NF190026, and W911NF2320006. This work was also supported in part by a subcontract awarded by the Pacific Northwest National Laboratory for the Secure Biosystems Design Science Focus Area “Persistence Control of Engineered Functions in Complex Soil Microbiomes” sponsored by the U.S. Department of Energy Office of Biological and Environmental Research.

## Author contributions

L. Y., Y. W. and E. Y. conceived the project and initial design of experiments. L. Y. and Y. W. designed and optimized new experimental methods, conducted all the experiments, analyzed data, and wrote the paper. W. L. assisted with data analysis and manuscript preparation. R. E. assisted with manuscript preparation. E. Y. provided overall supervision of the study, worked with R. E. to define project scope, and secured project funding for experiments, supplies, and equipment.

## Data availability

All data and code are available upon request.

## Method

### Plasmids construction

The plasmid used for the preparation of 20 kb DNA consists of a pBBR1-derived backbone carrying a kanamycin resistance marker and a synthesized insert fragment from Integrated DNA Technologies (IDT, USA). It also includes two buffer DNA fragments, which are derived from engineered *E. coli* RE1000^62^ and correspond to the upstream (genomic coordinates 282,501–287,037) and downstream (genomic coordinates 287,128–296,693) regions of the synthetic insert. The backbone was amplified from the pEYF1K plasmid^23^ using high-fidelity DNA polymerase (Vazyme, China, Cat. No. P520) with the following primers: Forward1 (5’-TCGTTCATAGAGTCCCGCTCAGAAGAACT-3’) and Reverse1 (5’-AGAGCATCAAACAGCGCACTTACGGGTT-3’). The two buffer DNA fragments were amplified from the genomic DNA of *E. coli* RE1000 using high-fidelity DNA polymerase (Vazyme, Cat. No. P520) and sequence-specific primers: Forward2 (5’-TGCGCTGTTTGATGCTCTATAACTTCCCGGC-3’), Reverse2 (5’-TAACTTGATGGGATGAGTGCCATGTGTTT-3’) for upstream buffer fragment and Forward3 (5’-ACTCATCCCATCAAGTTAGAGCGGTCCACC-3’), Reverse3 (5’-ACGTGCAGATGGGAACCGTGTAAATGCTCA-3’) for downstream buffer fragment. The synthesized insert fragment comprises a J23104 promoter-driven BFP gene, followed by a pTet-controlled tmBroccoli gene^40^, and a J23108 promoter-regulated RFP gene (Supplementary Material and Method). This insert was amplified by PCR using high-fidelity DNA polymerase (Vazyme, Cat. No. P520).

All four fragments were assembled using Gibson Assembly Master Mix (Vazyme, Cat. No. C117 or C116). The final plasmid construct, named pYW1, comprises a 3.7 kb backbone, a 4.7 kb upstream buffer fragment, a 1.9 kb insert, and a 10 kb downstream buffer fragment.

For the experiments in Figure 5, we replaced the original promoter and terminator region of the tmBroccoli gene in pYW1 with a T7 RNAP-specific promoter and a T7 terminator. This new DNA fragment was synthesized by IDT and amplified by PCR. The remaining backbone of the pYW1 plasmid was also prepared by PCR amplification. The two fragments were then assembled using Gibson assembly to create a new construct, named pYW1-T7.

### Linear DNA preparation

The 20 kb DNA fragment was amplified from plasmid pYW1 or pYW1-T7 using long-range PCR (Vazyme, Cat. No. PK512) with a biotin-modified forward primer (5’-CTTTGCTCAGTGTTCCGAATTCAATGCTC-3’) and a digoxigenin-modified reverse primer (5’-ACAATGTCATACCCGCTTTTTGACAAAGAC-3’). The PCR products were then separated by agarose gel electrophoresis using a 0.5% agarose (RPI, USA, Cat. No. A20090-50.0) gel prepared in Tris-acetate-EDTA (TAE) buffer to isolate the 20 kb target fragment from nonspecific products. The desired 20 kb band was isolated and purified using a DNA gel extraction kit (Vazyme, Cat. No. DC-301).

### Channel and DNA preparation for single-molecule assay

Commercial uncoated channel slides (ibidi, Germany, Cat. Nos. 80161, 80601, or 80621) were coated with 30 µg/mL anti-digoxigenin antibody (Bio-Rad Laboratories Inc., USA, Cat. No. 3210-0488) by incubating overnight at 4 °C. The slides were subsequently blocked overnight at 4 °C using blocking buffer composed of 10 mM potassium phosphate buffer (Sigma-Aldrich, USA, Cat. No. P0662, P3786), 10 mg/mL bovine serum albumin (BSA; Sigma-Aldrich, Cat. No. A7030), 0.22 mg/mL polyglutamic acid (Sigma-Aldrich, Cat. No. P4761), and 3 mM sodium azide (Sigma-Aldrich, Cat. No. S2002). After blocking, slides were washed twice with standard buffer containing 10 mM potassium phosphate buffer, mg/mL BSA, and 0.1% Tween-20 (Sigma-Aldrich, Cat. No. P9416). Slides were used immediately or stored at 4 °C for up to 1–2 days.

The 20 kb DNA fragment was amplified and purified as described above. For each channel, 500 ng of the purified DNA was labeled with Cyanine5 (Cy5) using the Label IT® Tracker Intracellular Nucleic Acid Localization Kit (Mirus Bio, USA; Cat. No. MIR3725) according to the manufacturer’s instructions. Excess dye was removed using a MicroSpin G-50 column (Cytiva Life Sciences, USA, Cat. No. 27533001).

The Cy5-labeled 20 kb DNA was immobilized on the pre-coated channel slide and interacted with streptavidin magnetic beads (New England Biolabs, USA, Cat. No. S1420S) via incubating the labeled DNA with the antibody-coated slide and magnetic beads for 1–2 hours at room temperature. Prior to use, magnetic beads were washed one with phosphate-buffered saline containing 1% BSA and then washed again with standard buffer supplemented with 1% BSA. After incubation, the slide was gently washed with standard buffer to remove unbound DNA and beads.

### Single-molecule transcription assay

Single-molecule DNA was mechanically stretched using a stack of magnets positioned at one side of the channel slide. For each channel, 30 µL of transcription buffer was prepared, consisting of 40 mM Tris-HCl (Cat. No. 10812846001), 150 mM KCl (Cat. No. P5405), 10 mM MgCl₂ (Cat. No. M4880), 1 mM DTT (Cat. No. 3860-OP), and 0.01% Triton® X-100 (Cat. No. T8787), all comes from Sigma-Aldrich. The transcription mixture was supplemented with 0.5 mM NTPs (NEB, Cat. No. N0466S) and 0.5 µL of *E. coli* RNA polymerase holoenzyme (NEB, Cat. No. M0551S).

For RNAP with topoisomerases groups, the transcription buffer was further supplemented with *E. coli* topoisomerase I (TOPO I; NEB, Cat. No. M0301S) and DNA gyrase (TopoGen Inc., USA, Cat. No. TG2000G) at a 0.2:1 ratio. To prepare the enzyme mixture, 1 µL of DNA gyrase was first diluted 5-fold in its dilution buffer. Then, 1 µL of diluted DNA gyrase combined with 1 µL of TOPO I to become a mixture. This mixture was then diluted 500-fold in the transcription buffer before use.

For experiments using DNA fragments with T7 promoter-driven tmBroccoli, transcription buffer was additionally supplemented with 0.5 µL of T7 RNA polymerase (NEB, M0251S).

When rifampicin was required, the transcription buffer was supplemented with 0.6 µL of a 50 µg/mL rifampicin solution to reach a final concentration of 1 µg/mL.

Immediately prior to imaging, the standard buffer in the channel slide was replaced with the transcription buffer respect to each group. DNA images were acquired every 2 minutes for a total of 15 timepoints.

### Co-localization of RNAP and single DNA molecule

For the co-localization assay, the transcription mixture was prepared in advance and incubated at room temperature for at least 1 hour prior to replacing the standard buffer in the channel slide. In addition to the components described above, the transcription mixture was supplemented with an anti-*E. coli* RNA polymerase primary antibody (BioLegend Inc., USA, Cat. No. 663903) at a 1:10,000 dilution, and an Alexa Fluor 594-conjugated secondary antibody against RNA polymerase at a 1:20,000 dilution (Invitrogen, USA, Cat.No. A-21145). For time-lapse imaging, DNA fluorescence images were acquired every 2 minutes for a total of 15 timepoints.

### DNA vibration analysis

DNA and transcription buffer were prepared as described above. For each timepoint, at least 30 consecutive frames of DNA images were captured as exposure time 200 milliseconds to analyze DNA vibration dynamics. Image acquisition was performed every 5 minutes over a total of 7 timepoints.

### Force extension analysis

Luer channel slides (ibidi, Germany; Cat. Nos. 80161 or 80621) were used for flow-based DNA extension analysis. DNA and transcription buffer were prepared as described above.

DNA molecules were extended by applying continuous buffer flow using a syringe pump. A range of flow rates, from 0 to 12 mL/hour, was applied to the channel. DNA was imaged and measured at each flow rate, and corresponding DNA lengths were recorded.

Bead displacements were also imaged at 20 milliseconds per frame, 350 frames in total to calculate the displacement variance at each flow rate.

Force is computed using the equation^33^ :

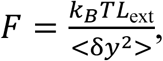

where *K*_B_ is the Boltzmann constant, *T* is the temperature, *L*_ext_ is the length of stretched DNA strand, and < δ*y*^2^ > is the variance of the fluctuations of the bead in the transverse direction, perpendicular to the optical axis. WLC model were used to fit the calculated and measured data, where contour length, *L_c_*, is 6.8 µm (20 kb DNA). The fitted WLC model is then used for force temporal dynamic analysis.

### Fluorescence microscopy

An Olympus IX81 TIR microscope, equipped with a 100x oil-immersion objective (CFI Apochromat TIRF 100XC Oil, NA 1.49, Nikon, Japan), a laser (Cobolt, Hubner-photonics), and a sCMOS camera (PrimeBSI, Teledyne Photometrics, Canada), was used to image Cy5 labeled DNA, RNAP–Alexa594.

### *In vitro* transcription assay

The transcription level of the tmBroccoli gene was analyzed using an *in vitro* transcription assay. Reactions were carried out in a transcription buffer supplemented with 2.5 mM NTPs, mM DFHBI 1T^40^ (R&D systems Inc., USA, Cat. No. 5610), and 1 µL of *E. coli* RNA polymerase holoenzyme. Each reaction had a total volume of 15 µL and contained 14 pmol of the prepared 20 kb DNA fragment as the transcription template. For selected experimental groups, the RNA polymerase inhibitor rifampicin was added at a final concentration of 1 µg/mL (Sigma-Aldrich, Cat. No. R3507), or varying amounts of *E. coli* topoisomerase I (NEB, Cat. No. M0301S) and DNA gyrase (TopoGen Inc., Cat. No. TG2000G) were included as indicated. Fluorescence from tmBroccoli-bound DFHBI 1T was measured using a 384-well microplate reader with an excitation wavelength of 472 nm and an emission wavelength of 507 nm. Readings were recorded every 2 minutes over a total of 721 cycles.

### Image analysis and peak detection

Time-lapse images from both fluorescence and bright-field channels were analyzed using Fiji software. Individual DNA molecules were cropped with the Fiji ROI manager along each timepoint. The long axis of each stretched DNA strand was defined as the x-axis. For each timepoint, a kymograph was generated along the x-axis to visualize spatial-temporal changes in DNA fluorescence intensity. To quantify fluorescence distribution, the mean intensity along the y-axis was calculated across the x-axis and plotted. These intensity profiles were smoothed using a Savitzky–Golay filter to aid peak detection. Peaks corresponding to DNA plectonemes or RNA polymerase (RNAP) were identified using the scipy.find_peaks algorithm (SciPy library, Python). Peak width was defined as the distance between the two intersection points of the horizontal line where its y-value is defined as the closest inflation point detected on either side of the peak and the smoothed mean intensity line. To analyze temporal change in the length of DNA strand, relative DNA length is expressed as a percentage, with the first timepoint for each DNA molecule set as 100% and subsequent timepoints calculated relative to this reference. To analyze the temporal dynamics of plectoneme position and size, all DNA strands lengths were normalized to a [0, 1] scale, where 0 and 1 represents the digoxigenin-modified end and biotin-modified end of the DNA strand, respectively. Relative peak position and relative peak size were calculated and compared using this normalized axis. Peak position distributions were computed for each frame using Gaussian KDE (Silverman’s bandwidth method). For frames containing less then 2 peaks, null distribution was assigned zero density. For each frame, density was normalized to half of its maximum density value for visual presentation. For vibration image analysis, the variance of the averaged position of the top 10% most intensity pixel on the y-axis (width of DNA) over 30 frames are calculated for each x-position (length of DNA).

### Statistical analysis

For comparisons between two groups, statistical significance was assessed using an unpaired, two-tailed t-test. For comparisons among multiple time-series groups, mixed-effects models were performed, followed by Tukey’s Honestly Significant Difference (HSD) post-hoc test. A p-value of less than 0.05 was considered statistically significant. For relative peak size and timepoints correlation analysis, a weighted Pearson’s correlation analysis was employed. Observations were weighted by a composite score prioritizing IQR (90%) and whisker range (10%).

