## Supplementary information for "Single-Molecule Imaging Reveals Transcription-Driven Supercoiling in Unconstrained DNA"

**Lili Yang[1,*], Yanran Wang[2,*], Wei Lyu[3], Robert. G. Egbert[4] and Enoch Yeung[1,2]**

[1]Institute for Collaborative Biotechnologies, University of California, Santa Barbara, CA 93106
[2]Department of Mechanical Engineering, University of California, Santa Barbara, CA 93106
[3]Department of Mathematics and College of Creative Studies, University of California, Santa Barbara, CA 93106
[4]Earth and Biological Sciences Division, Pacific Northwest National Laboratory, Richland, WA 99354
[*]Those authors contribute equally.

**Correspondence:**
Enoch Yeung

(805) 893-4087
3104 BioEngineering, University of California, Santa Barbara,
Santa Barbara, CA 93106-5070

#### Supplementary Figures

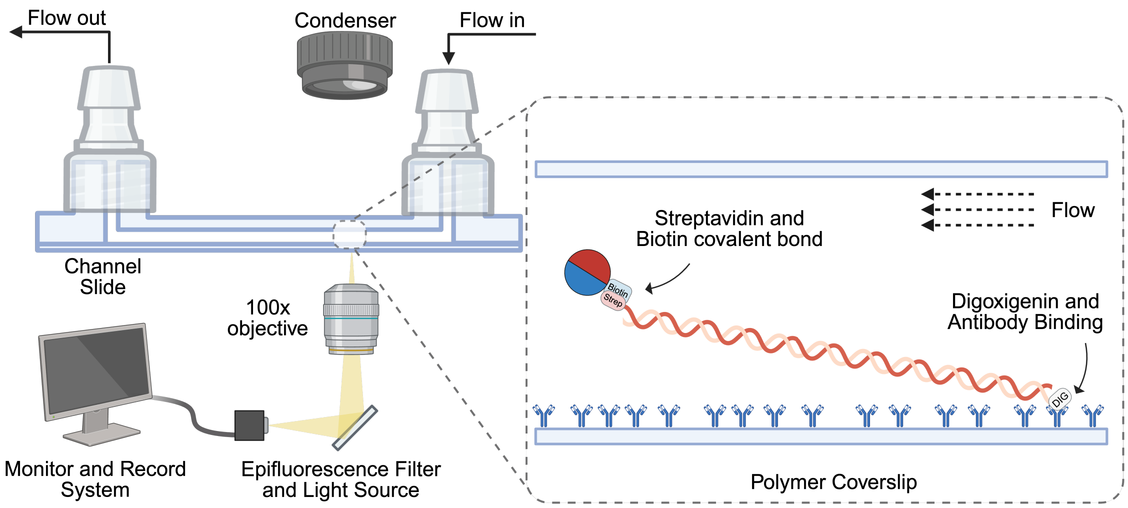


**Figure S1. Schematic of the single-DNA-molecule stretching experiment using flow.** A 20-kb DNA molecule is asymmetrically labeled with digoxigenin (DIG) at the 5' end of one strand and biotin at the 5' end of the complementary strand. The DIG-labeled end is tethered to the surface of a $\mu$-Slide via an anti-DIG antibody, while the biotinylated end is attached to a streptavidin-coated paramagnetic bead. A constant flow from a syringe pump stretches the DNA horizontally. DNA conformations are monitored in real time using an inverted fluorescence microscope. This figure was created in BioRender (https://BioRender.com).

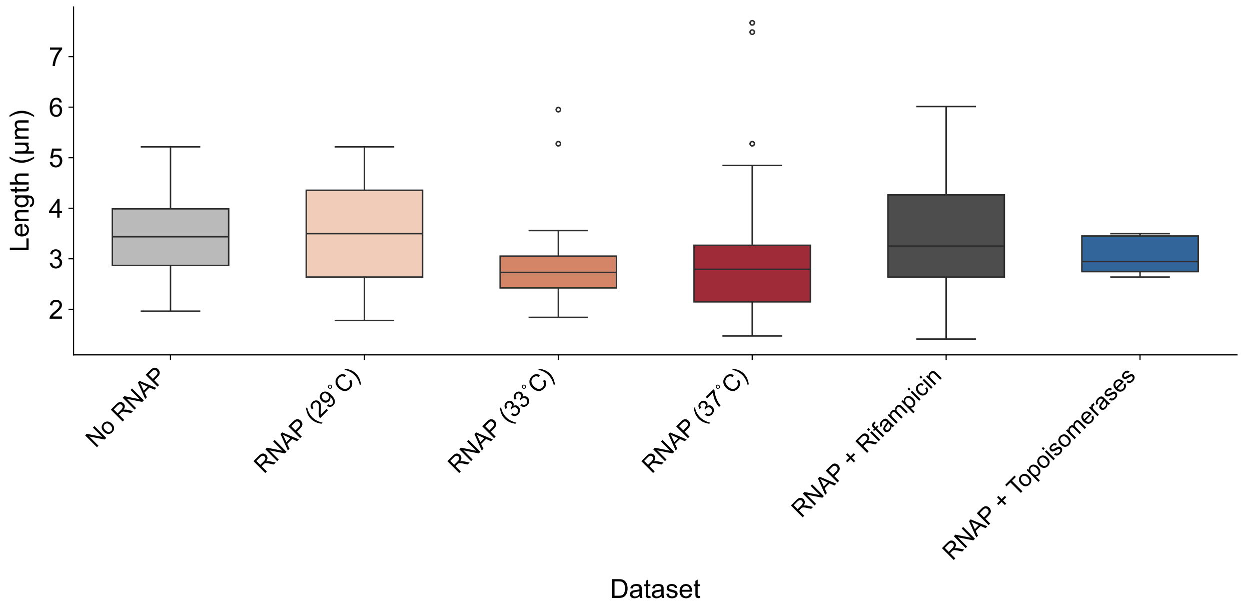


**Figure S2. Initial DNA length in each experimental condition.** DNA lengths were calculated from the initial observation time point for each group. The No RNAP, RNAP + Rifampicin, and RNAP + Topoisomerases groups were measured at 37°C. The RNAP-only condition was measured at 29°C, 33°C, and 37°C. Conditions are defined as: **No RNAP**, DNA incubated with transcription buffer only; **RNAP + Rifampicin**, transcription buffer supplemented with the *E.coli* RNAP inhibitor rifampicin at 1 $\mu$g/mL; **RNAP + Topoisomerases**, transcription buffer supplemented with a combination of gyrase and topoisomerase I at a 0.2:1 ratio. DNA molecules in each group: No RNAP (N = 48), RNAP at 29°C (N = 17), RNAP at 33°C (N = 26), RNAP at 37°C (N = 60), RNAP + Rifampicin (N = 47), RNAP + Topoisomerases (N = 8).

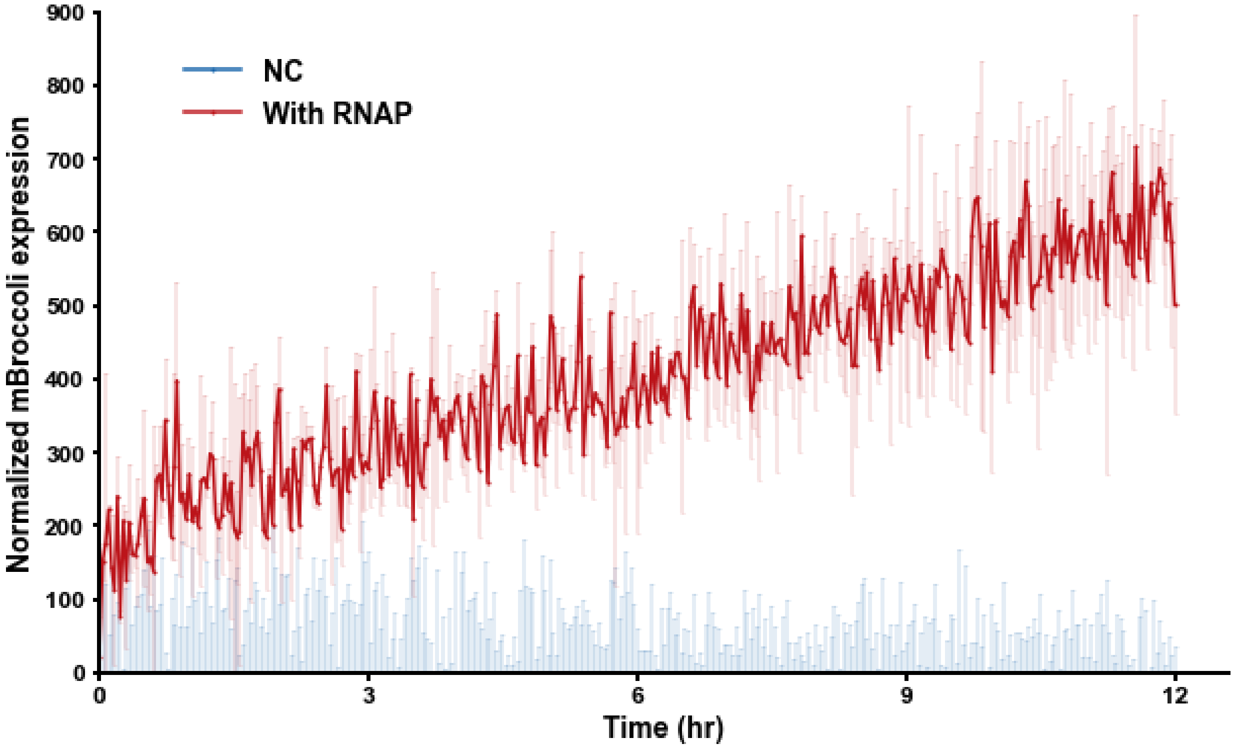


**Figure S3. Time series transcription with a Cy5-labeled DNA template.** A PCR-amplified 20-kb DNA template was gel-extracted, labeled with Cy5, and used to monitor tmBroccoli expression over 12 hours in a plate reader. Each 15 $\mu$L reaction contained 14 nM of the DNA template. The tmBroccoli fluorophore, DFHBI, was included in the transcription buffer, and fluorescence was measured every 2 minutes. Signals were normalized to negative controls (NC) containing all reaction components except DNA. Shaded regions represent the standard deviation.

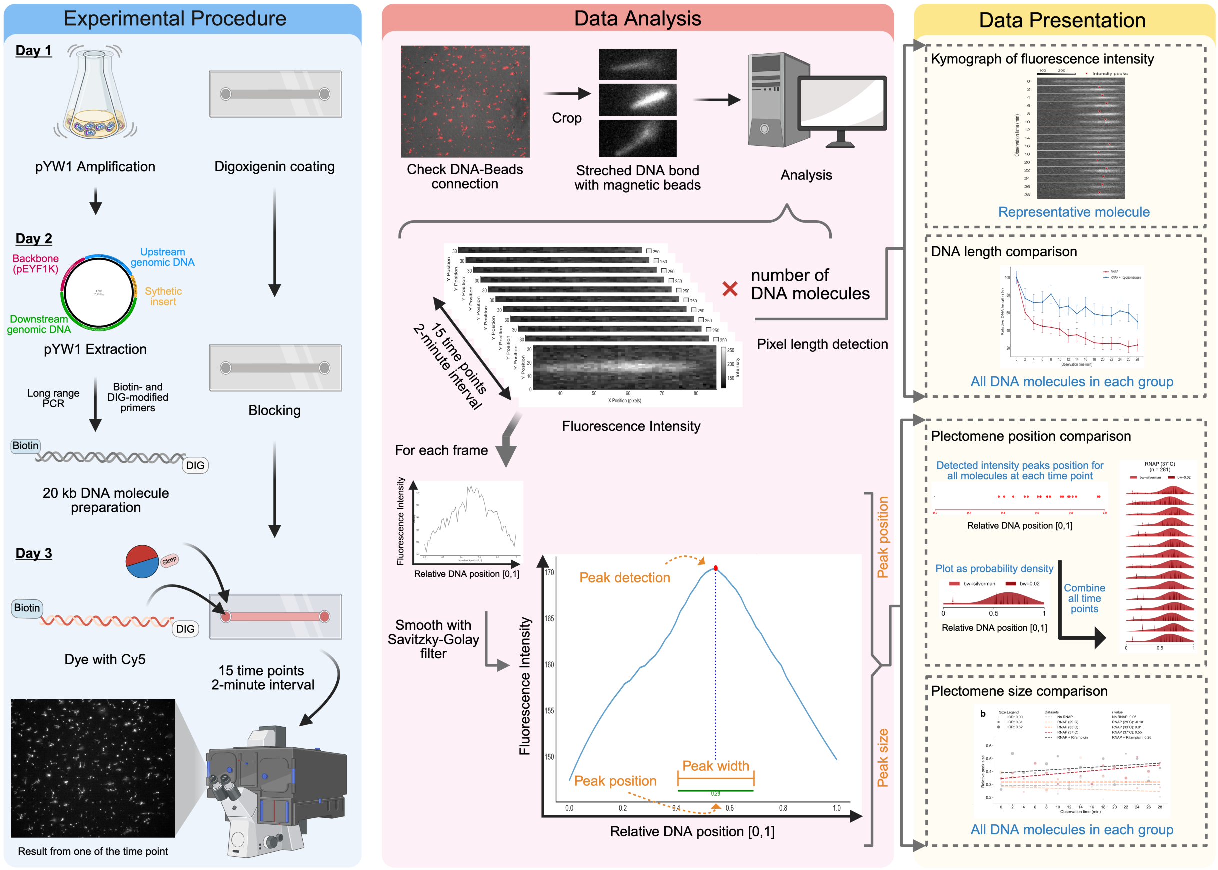


**Figure S4. Schematic overview of the experimental and analytical workflow.** The schematic outlines the overall data processing pipeline, from sample preparation to final interpretation.This illustration figure was created in BioRender.

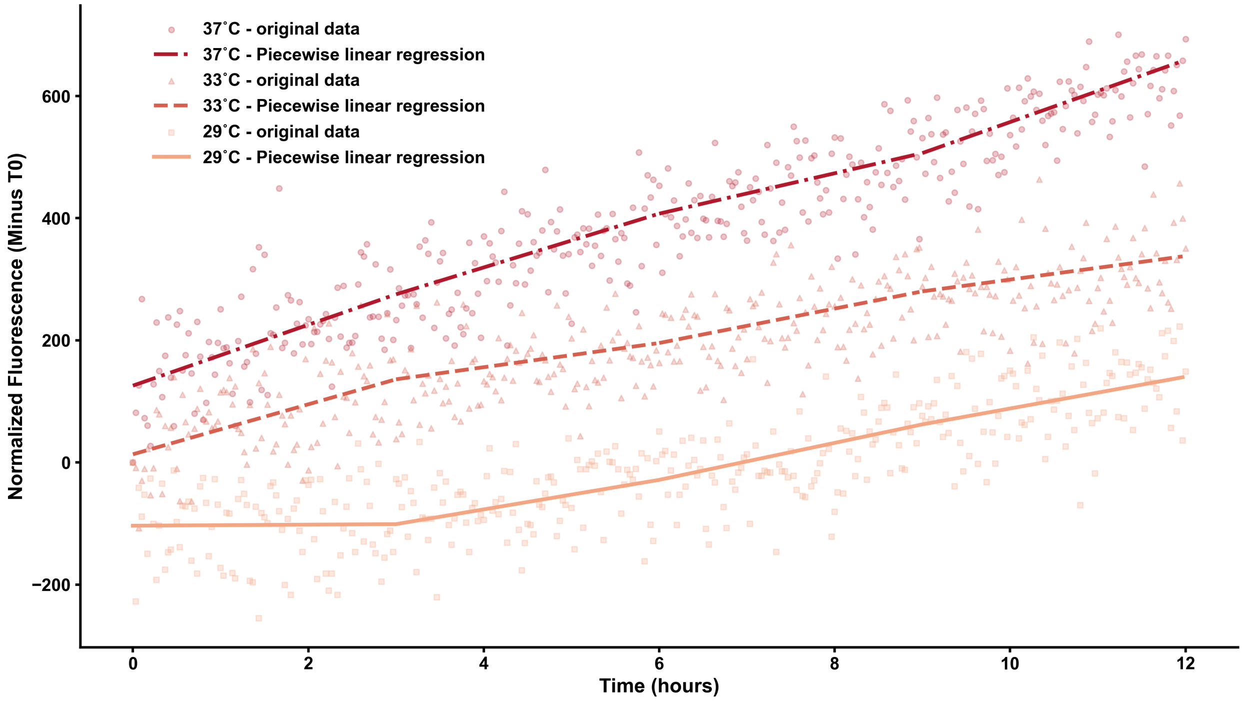


**Figure S5. Temperature modulates the RNAP transcription rate on unconstrained linear DNA.** A 20-kb linear DNA template was amplified from pYW1 by PCR and purified by gel extraction. DHFBI was added to all groups. The tmBroccoli signal was measured every 2 minutes. Fluorescent signals were normalized to respective negative controls, which contained all elements except DNA.

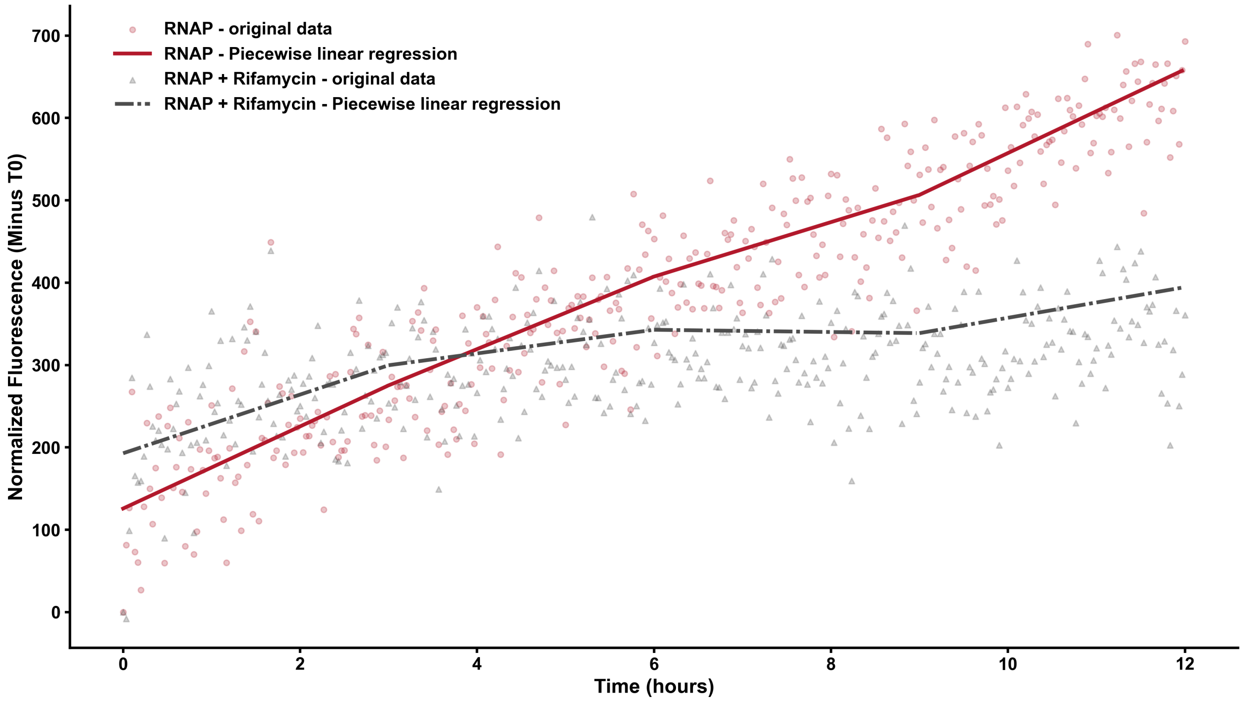


**Figure S6. Rifampicin inhibits RNAP transcription rate on unconstrained linear DNA.** A 20-kb linear DNA template was amplified from pYW1 by PCR and purified by gel extraction. Transcription by *E.coli* RNAP was suppressed using the inhibitor rifampicin at a final concentration of 1 $\mu$g/mL. DFHBI was added to all groups. The tmBroccoli fluorescence signal was measured every 2 minutes and normalized to that of negative controls, which contained all reaction components except DNA.

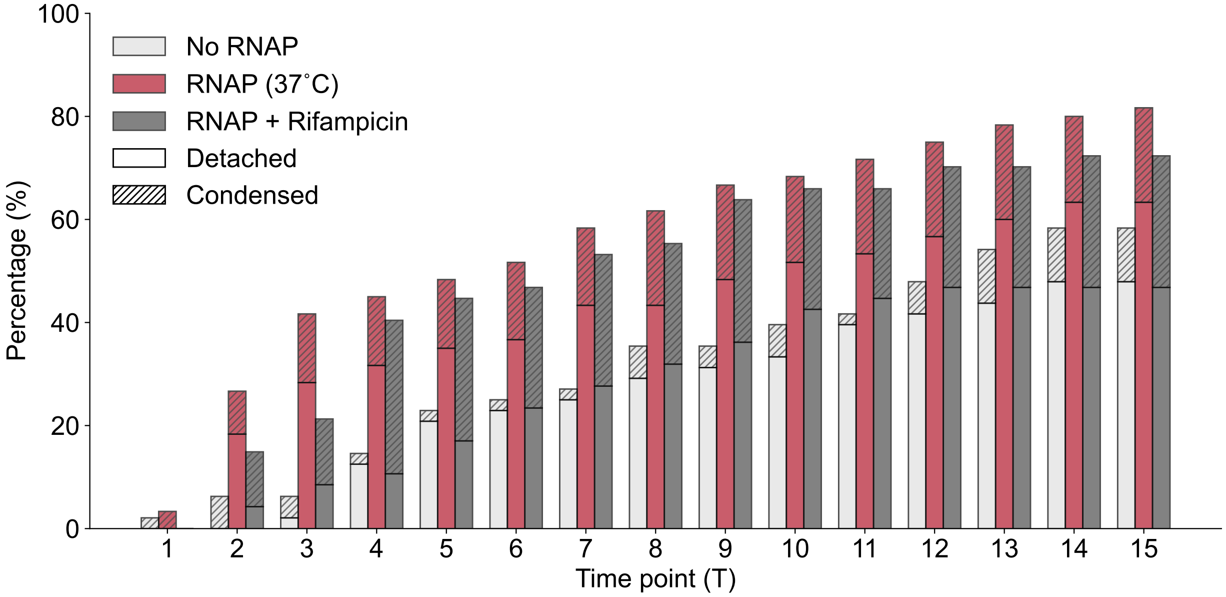


**Figure S7. DNA state transitions over time.** The state of individual DNA molecules was recorded at 2-minute intervals. States are defined as follows: **Detached**, molecules that disappeared from the microscope field; **Condensed**, molecules compacted into a single spot (size <1.2 µm). All experiments were conducted at 37°C. In the RNAP + Rifampicin group, the *E.coli* RNAP inhibitor was added to the transcription buffer at a final concentration of 1 $\mu$g/mL. Sample sizes: RNAP at 29°C (n = 80), RNAP at 33°C (n = 199), and RNAP at 37°C (n = 281).

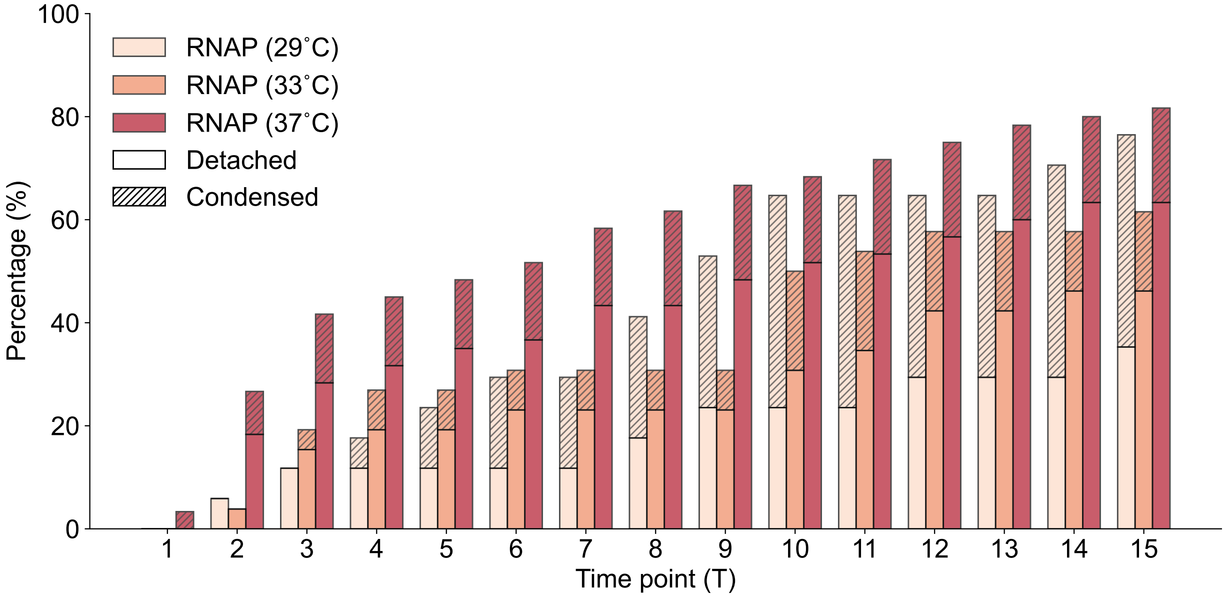


**Figure S8. Attenuated transition of DNA state at lower temperature.** The state of individual DNA molecules was recorded at 2-minute intervals. States are defined as follows: **Detached**, molecules that disappeared from the microscope field; **Condensed**, molecules compacted into a single spot (size <1.2 µm). Experiments were conducted at 29°C, 33°C, and 37°C. Sample sizes: No RNAP (n = 362), RNAP at 37°C (n = 281), and RNAP + Rifampicin (n = 265).

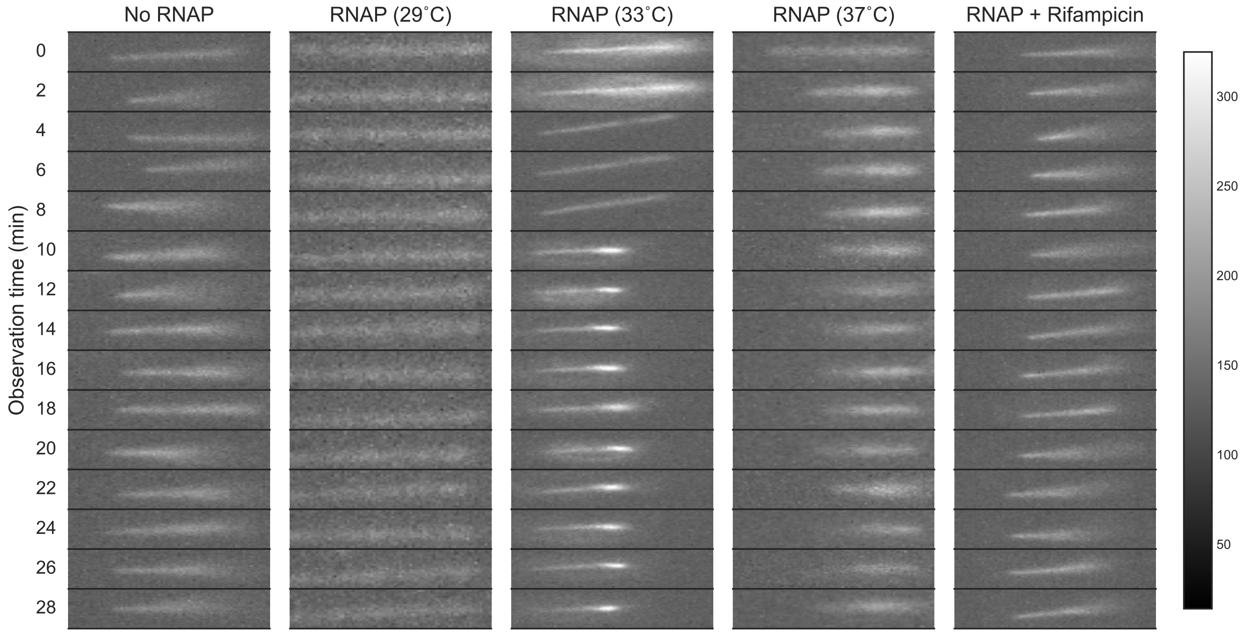


**Figure S9. Representative kymograph of real-time analysis of DNA conformational changes.** All groups were performed in the presence of transcription buffer and *E.coli* RNAP at different temperature or with 1$\mu$g/mL of *E.coli* RNAP inhibitor rifampicin.

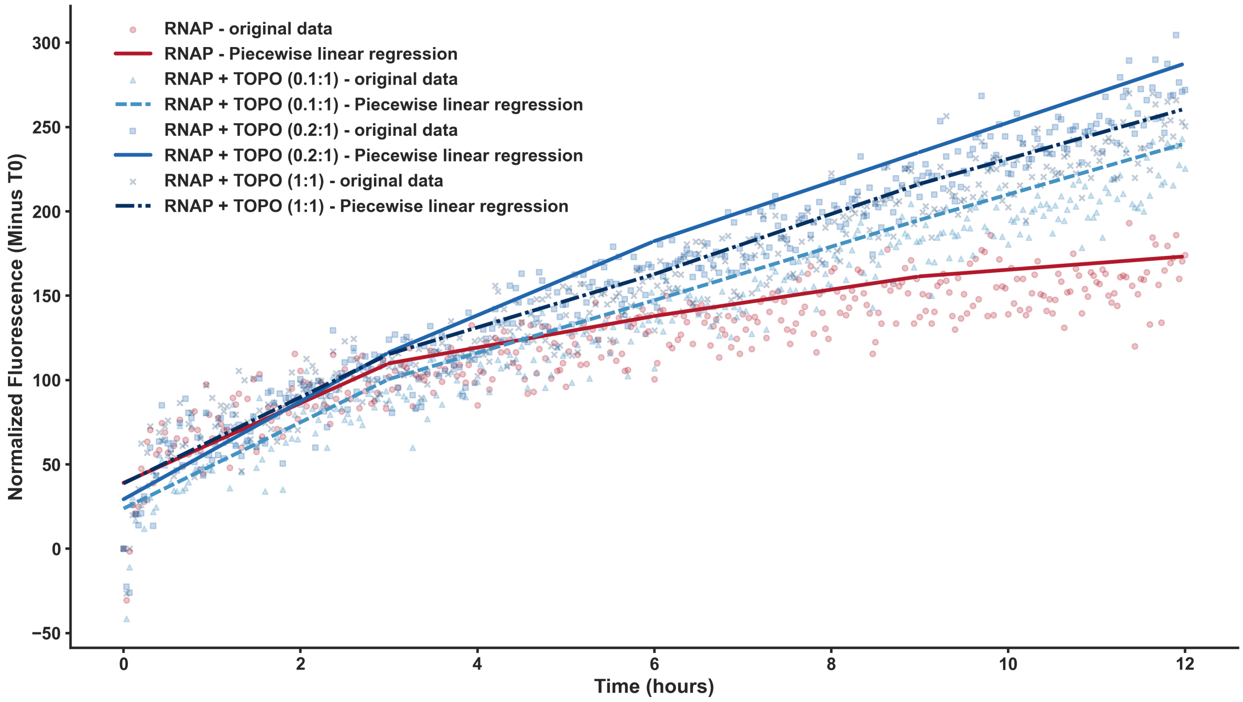


**Figure S10. Topoisomerases accelerate RNAP transcription rate on unconstrained linear DNA.** A 20-kb linear DNA template was amplified from pYW1 by PCR and purified by gel extraction. DHFBI was added to all groups. The tmBroccoli signal was measured every 2 minutes. Fluorescent signals were normalized to their respective negative controls, which contained all elements except RNAP. TOPO (X : Y) refers to the ratio of reaction units between gyrase and topoisomerase I.

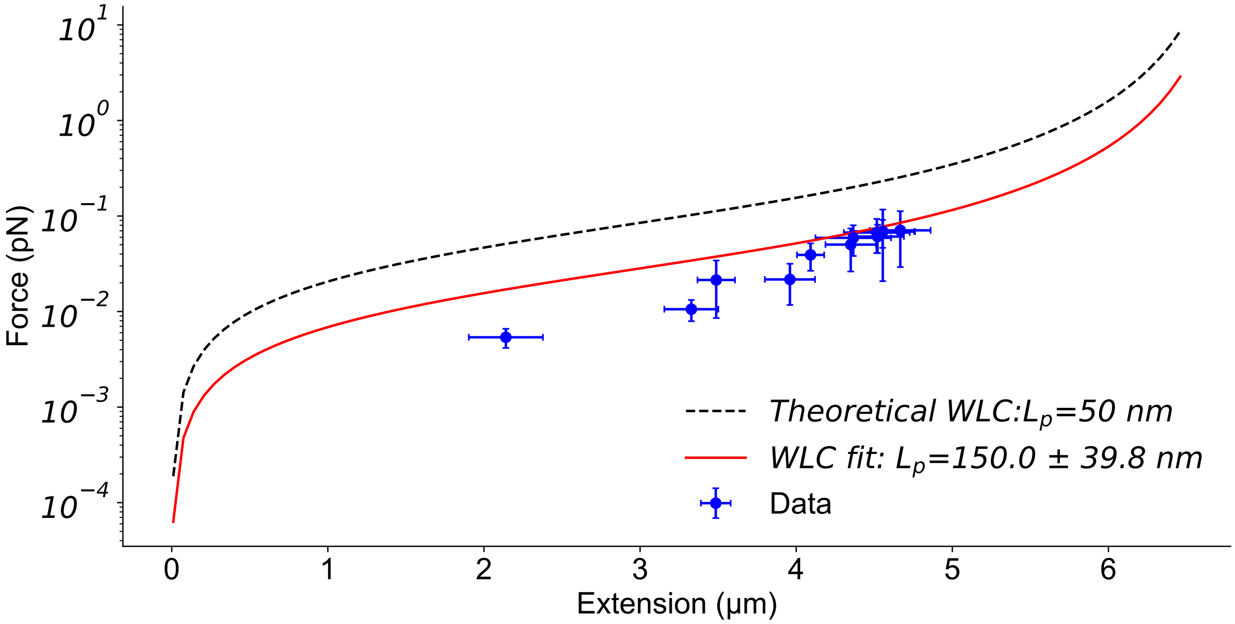


**Figure S11. Force vs. Extension curve fit.** The Marko-Siggia worm-like chain (WLC) approximation was fitted to the data using a constant $L_c$ value. Error bars represent the average standard deviation ($n$ = 5).

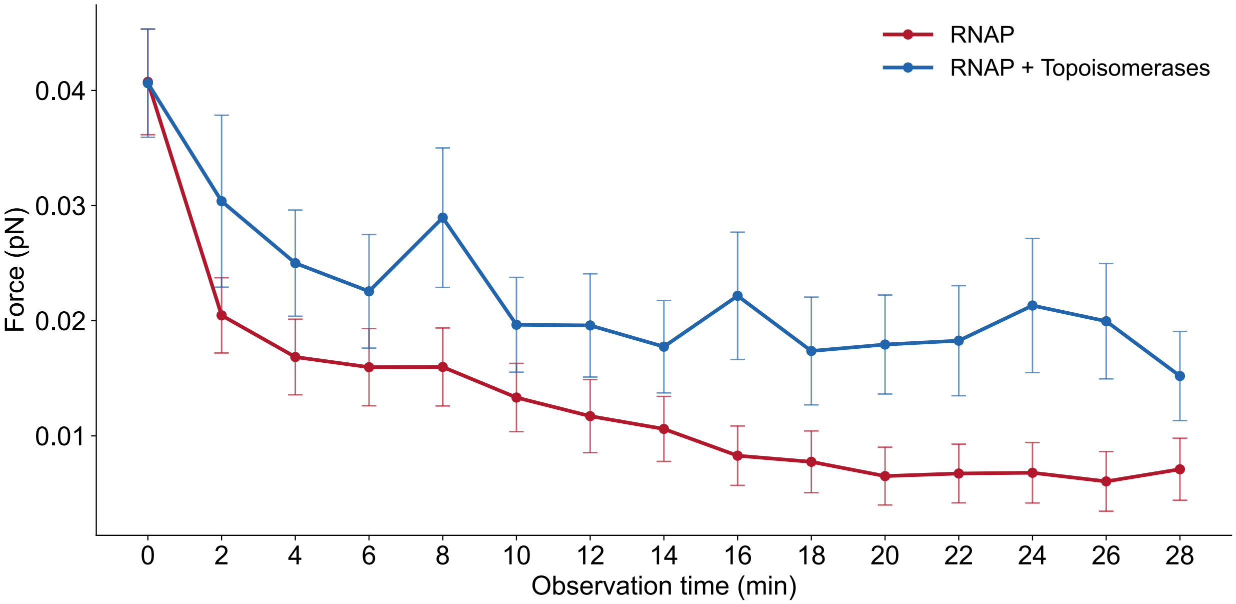


**Figure S12. Topoisomerases modulate transcription-induced DNA compaction force generated by RNAP.** Force was calculated using the Marko-Siggia worm-like chain (WLC) approximation with a fitted $L_p$ value. DNA lengths were gathered from the RNAP group and the RNAP + Topoisomerases group in Figure 3d. In the RNAP + Topoisomerases group, the transcription buffer was supplemented with a combination of gyrase and topoisomerase I at a 0.2:1 ratio. Data are presented as mean ± standard deviation.

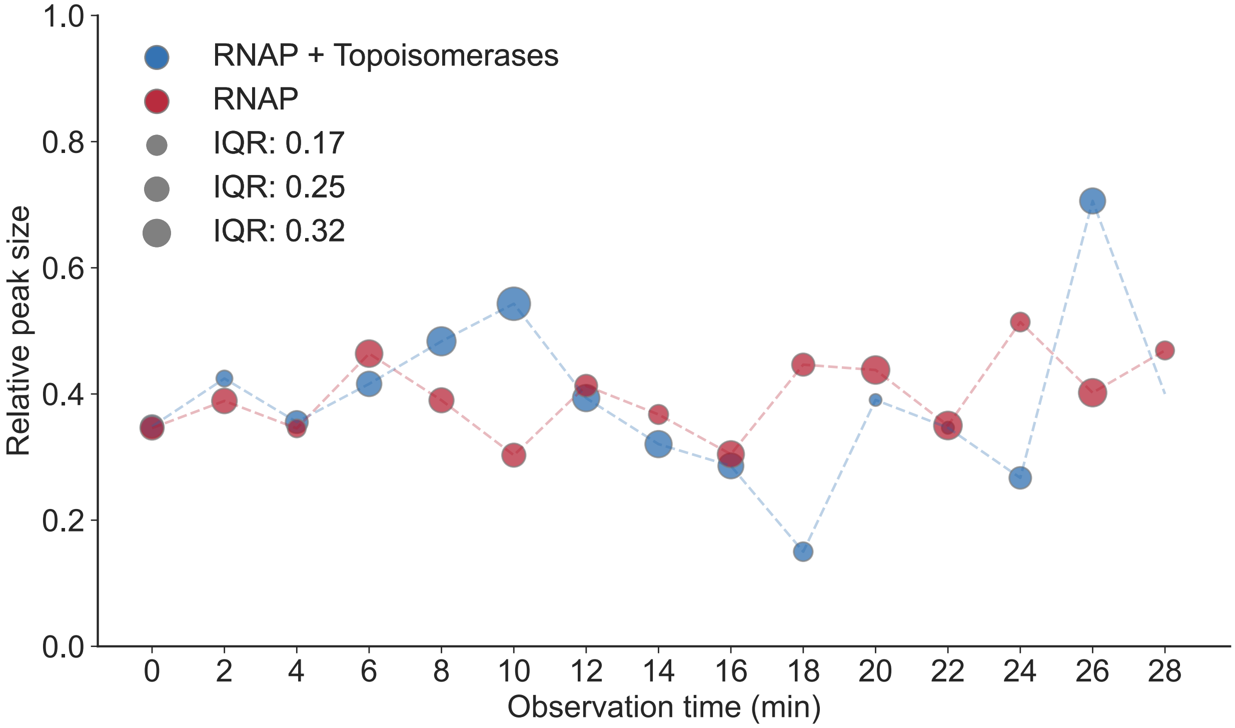


**Figure S13. Topoisomerases modulate plectoneme size generated by RNAP.** DNA peak size normalized to DNA length over time. The interquartile range (IQR) is shown for each time point.

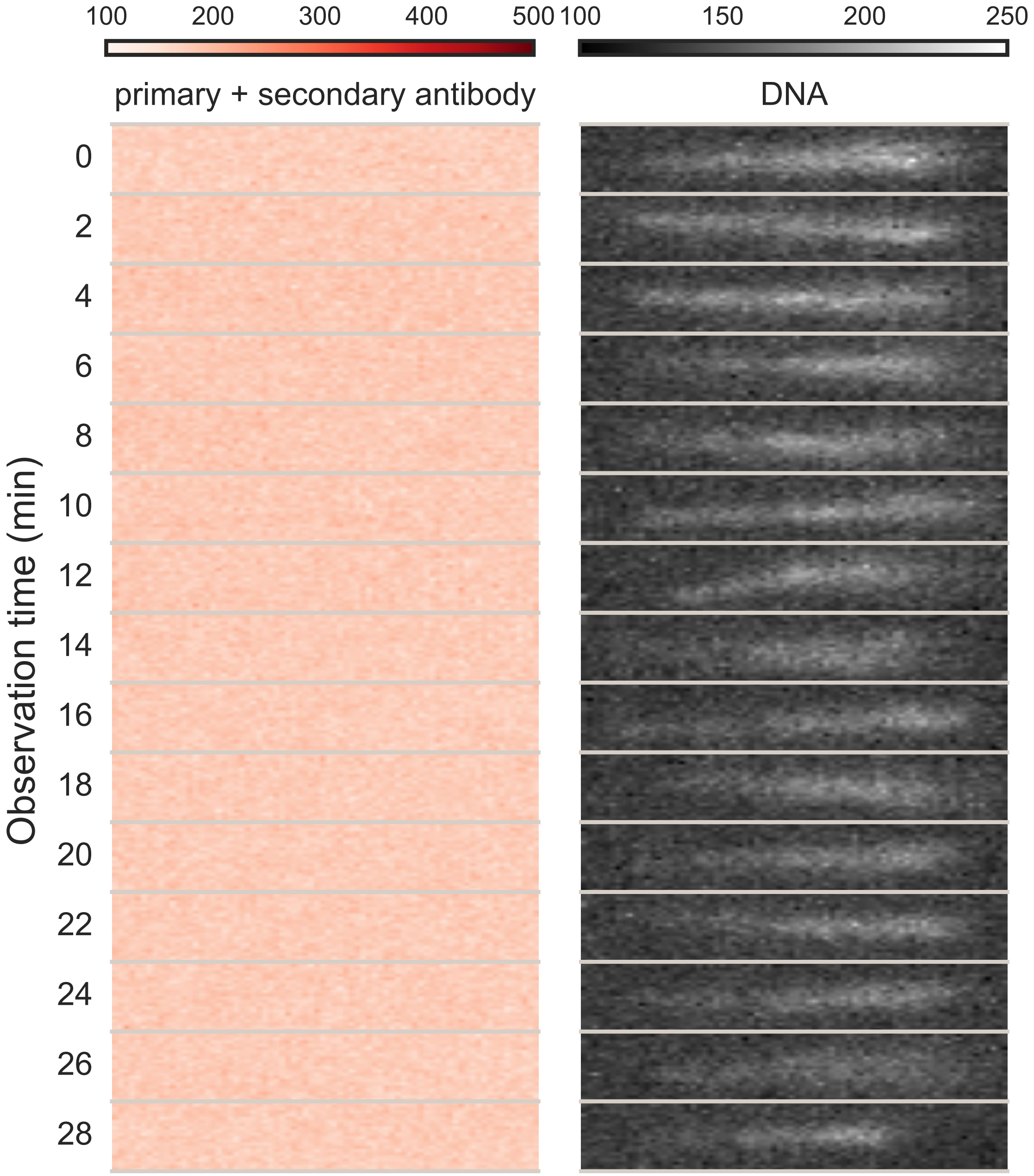


**Figure S14. Alexa 594-conjugated secondary antibody and anti-*E. coli* RNAP primary antibody exhibits no non-specific binding to DNA.** Real-time analysis of antibodies–DNA co-localization dynamics, shown as a time-series image: Alexa 594-conjugated sencondary antibody with primary antibody targeting RNAP(left) and Cy5-labeled DNA (right). The time interval between frames is 2 minutes.

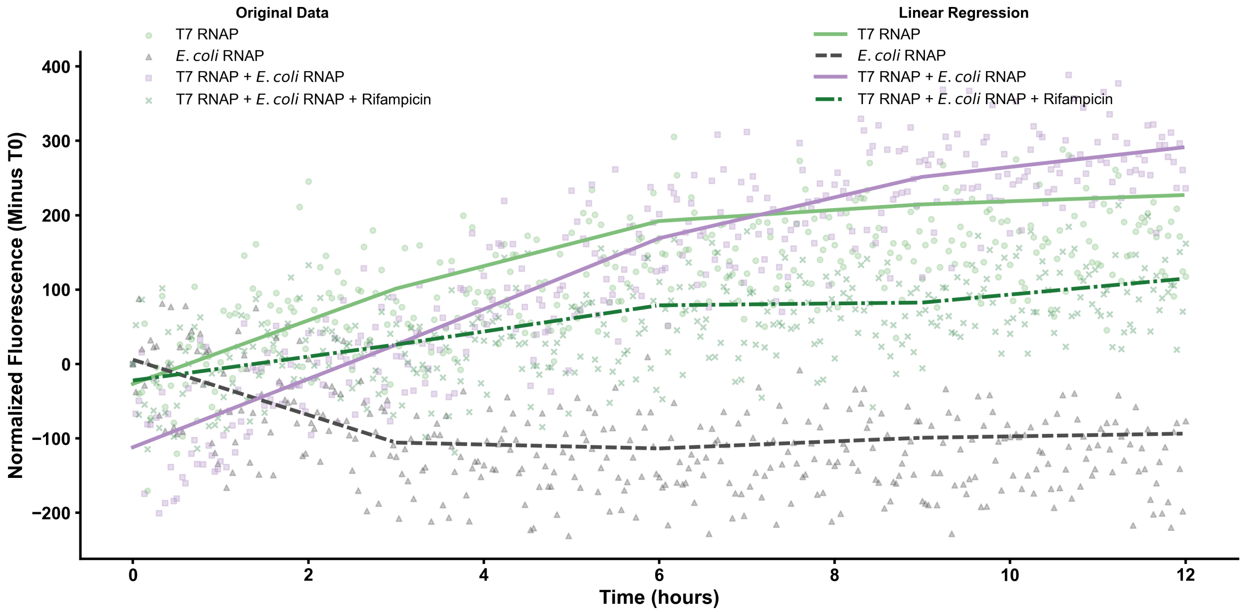


**Figure S15. T7 RNAP drives single gene expression.** A 20-kb DNA template was amplified from pYW1-T7, gel-extracted, and dyed with Cy5. This template was used to monitor tmBroccoli expression over a time series of up to 12 hours in a plate reader. Each 15 $\mu$L reaction contained 14 nM of the DNA template. The transcription buffer was supplemented with the tmBroccoli fluorophore, DHFBI. Fluorescence was measured every 2 minutes. Signals were normalized to negative controls containing all reaction components except DNA in the T7 RNAP + $E.coli$ RNAP + Rifampicin group. In the rifampicin-treated group, the antibiotic was added to the transcription buffer at a final concentration of 1 $\mu$g/mL.

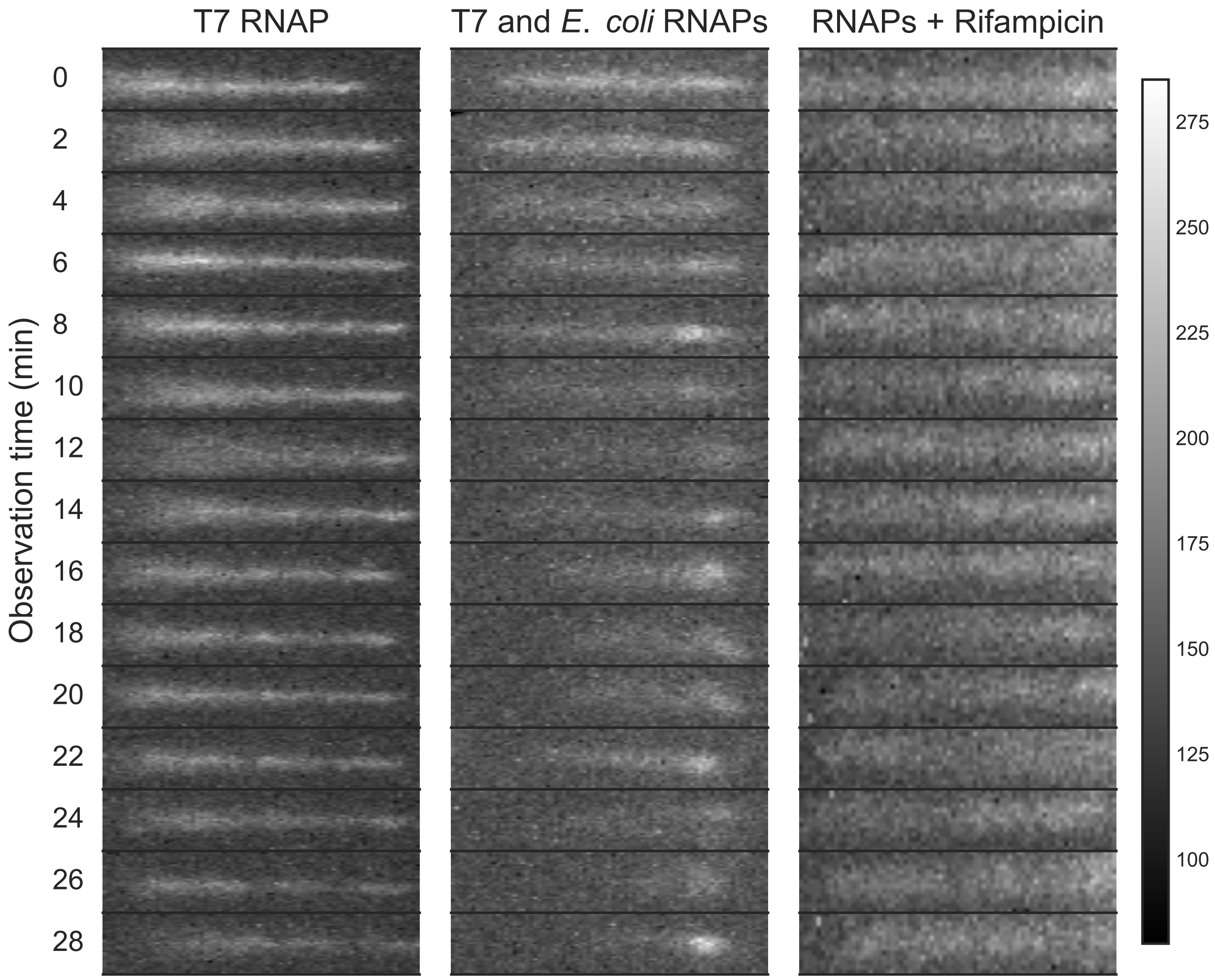


**Figure S16. Representative kymograph of real-time analysis of DNA conformational changes with combination of T7 RNAP and *E. coli* RNAP.** All groups were performed in the presence of transcription buffer and T7 RNAP without or with *E.coli* RNAP. Final concentration of *E.coli* RNAP inhibitor rifampicin is 1$\mu$g/mL.

### Supplementary Material and Method

#### Sequence of insert genes

1 ATCAAGTTAGAGCGGTCCACCCGTTTTGACAGCTAGCTCAGTCCTAGGTATTGTGCTAGC

61 TCTAGAGTCACACAGGAAAGTACTAGATGAGTGAGCTTATTAAAGAGAACATGCACATGA

121 AGTTGTATATGGAAGGGACCGTCGACAATCATCATTTCAAGTGCACTTCTGAAGGTGAAG

181 GGAAGCCGTATGAGGGAACCCAAACTATGCGTATTAAAGTAGTCGAAGGAGGGCCTTTGC

241 CATTCGCCTTTGATATTCTGGCGACAAGTTTTTATATGGATCTAAGACTTTTATCAACC

301 ACACGCAAGGGATCCCGGATTTCTTTAAGCAATCCTTCCCCGAAGGCTTTACATGGGAAC

361 GCGTGACAACATACGAGGACGGGGGCGTCTTGACTGCTACACAAGATACTAGTTTGCAAG

421 ATGGTTGTCTTATCTATAATGTTAAAATCCGTGGAGTGAACTTTACTAGTAACGGTCCGG

481 TCATGCAAAAGAAAACGCTGGGGTGGGAAGCCTTCACGGAAACCCTGTATCCCGCGGATG

541 GCGGTTTGGAGGGGCGCAACGATATGGCGTTGAAGTTAGTTGGTGGCTCACACCTTATCG

601 CCAATATTAAGACCACGTATCGCTCGAAGAAACCAGCTAAAAACTTAAAGATGCCGGGCG

661 TGTACTACGTGGACTACCGCCTTGAGCGCATCAAGGAAGCAAATAACGAAACTTATGTCG

721 AGCAGCATGAGGTAGCGGTCGCCCGCTATTGCGATCTTCCAAGTAAACTTGGCCATAAAC

781 TGAACTGACGCAAAAAACCCCGCTTCGGCGGGGTTTTTTCGCGAGCTGTTGACAACTCTA

841 TCATTGATAGAGTTATAATGTTCCCTATCAGTGATAGAGACGCTTGCCATGTGTATGTGG

901 GAGACGGTCGGGTCCAGATATTCGTATCTGTCGAGTAGAGTGTGGGCTCCCACATACTCT

961 GATGATCCTTCGGGATCATTCATGGCAATTCAGCCAAAAAACTTAAGACCGCCGGTCTTG

1021 TCCACTACCTTGCAGTAATGCGGTGGACAGGATCGGCGGTTTTCTTTTCTCTTCTCAACT

1081 GACAGCTAGCTCAGTCCTAGGTATAATGCTAGCTCTAGAGAAAGAGGGGAAATACTAGAT

1141 GGCTTCTAGCGAGGACGTTATTAAAGAATTCATGCGCTTTAAGGTTCGTATGGAGGGTTC

1201 AGTAAATGGCCACGAATTCGAAATCGAGGGTGAGGGAGAAGGCCGCCCGTATGAAGGGAC

1261 TCAAACGGCCAAGTTGAAGGTGACAAAGGGGGGACCGTTGCCTTTCGCGTGGGATATCCT

1321 TTCGCCTCAATTTCAATATGGGTCAAAAGCGTATGTCAAGCACCCAGCCGATATTCCGGA

1381 TTACTTAAAACTTAGTTTCCCGGAGGGTTTCAAGTGGGAGCGCGTTATGAATTTTGAGGA

1441 CGGGGGAGTTGTTACGGTAACGCAAGACAGCTCTCTGCAGGATGGGGAGTTTATCTATAA

1501 AGTAAAACTTCGTGGAACTAACTTCCCTTCAGACGGCCCAGTCATGCAGAAGAAGACTAT

1561 GGGTTGGGAAGCCAGCACTGAGCGCATGTATCCGGAGGACGGTGCATTGAAGGGAGAGAT

1621 CAAGATGCGCTTAAAATTGAAAGACGGAGGGCATTATGATGCTGAAGTCAAAACGACTTA

1681 TATGGCAAAGAAACCGGTACAGTTACCGGGGGCCTATAAGACAGACATCAAACTTGATAT

1741 CACCTCCCACAACGAAGACTACACCATTGTGGAACAGTATGAGCGTGCCGAAGGTCGCCA

1801 TTCAACCGGAGCATAACGCCAAAAACCCCGCTTCGGCGGGGTTTTTCCGCGCAGTGAGCA

1861 TTTACACGGTTCCCA

**Legend:**

- Red: RFP region;
- Blue: BFP Region;
- Green: tmBroccoli aptamer with F30 scaffold;
- Yellow: J23104 promoter;
- Orange: pTet promoter;
- Purple: J23108 promoter.
